# The distribution of fitness effects of new mutations in regulatory regions of the *D. melanogaster* genome

**DOI:** 10.64898/2026.03.01.708907

**Authors:** Austin Daigle, Jacob Marsh, Andrew Kay, Parul Johri

## Abstract

Although non-coding regions play important roles in gene regulation and contribute to individual fitness, the precise distribution of fitness effects (DFE) of new mutations in these regions remains poorly understood. Here, we carefully compile experimentally validated regulatory regions in non-coding regions in the *Drosophila melanogaster* genome and identify putatively neutral sites near them. Incorporating a realistic genomic architecture that mimics the placement of the regulatory and coding regions, as well as a realistic heterogeneity in recombination and mutation rates across the genome, we use forward-in-time simulations to assess the power and accuracy of population genetics approaches that infer the DFE of new mutations in these regions. While the parameters of DFEs primarily comprising moderately and strongly deleterious mutations are estimated accurately, those of DFEs comprising mostly mildly deleterious mutations are misinferred. Applying these insights to three African *D. melanogaster* populations, we find that a large fraction of new mutations in functionally important non-coding regions are moderately deleterious, as opposed to strongly deleterious in coding regions. While the fraction of beneficial substitutions in regulatory regions (0.25-0.45) was also lower than in coding regions (∼0.5), our results suggest that non-coding regions contribute a majority of new deleterious mutations and beneficial substitutions in *D. melanogaster* populations. By incorporating both the genomic distribution and the inferred DFE of non-coding regions, we demonstrate that the effects of background selection across the genome are more accurately captured than with coding regions alone, highlighting the importance of considering selection on non-coding regions when interpreting patterns of genomic variation.

**AUTHOR SUMMARY:** The distribution of fitness effects (DFE) of new mutations in non-coding regulatory regions remains poorly characterized, despite the importance of these regions in gene regulation and evolution. Here, we investigate the DFE of mutations in regulatory regions in the *Drosophila melanogaster* genome by combining functional genomic annotations with population genetic inference. Using forward-in-time simulations incorporating realistic genomic architecture and variation in mutation and recombination rates, we evaluated the accuracy of three commonly used DFE inference methods. We found that DFEs skewed towards moderately and strongly deleterious mutations could be inferred reliably, whereas inference of DFEs skewed towards weakly deleterious mutations was more sensitive to positive selection. Applying these approaches to three African *D. melanogaster* populations, we inferred that mutations in conserved and experimentally validated regulatory regions are generally less deleterious than nonsynonymous coding mutations. Nevertheless, because non-coding regions comprise a substantially larger fraction of the genome, our results indicate they likely contribute the majority of new deleterious mutations and beneficial substitutions. Furthermore, incorporating selection on non-coding regions improved predictions of background selection across the genome. Our findings demonstrate the importance of selection on regulatory regions in shaping genetic diversity.

## INTRODUCTION

The distribution of fitness effects (DFE) of new mutations is a central quantity in population genetics because it measures how mutations impact fitness, governs the maintenance of genetic variation, and shapes genome-wide patterns of polymorphism and divergence. Accurate inference of the DFE is necessary to estimate the genomic burden of segregating deleterious alleles (Eyre-Walker and Keightley 2007; Agrawal and Whitlock 2012; Henn et al. 2015), the prevalence of beneficial substitutions (Smith and Eyre-Walker 2002; Eyre-Walker and Keightley 2009; Messer and Petrov 2013), and the magnitude and spatial distribution of background selection across genomes (Nordborg et al. 1996; Johri et al. 2020; Charlesworth and Jensen 2021).

Studies of the DFE have typically focused on nonsynonymous mutations in protein-coding regions. Population-genetic approaches to infer the DFE usually follow a two-step strategy using the site-frequency spectrum (SFS) and divergence data (reviewed in Johri et al. 2022). First, patterns at putatively neutral sites (e.g., synonymous or short intronic sites) are used to characterize how the observed SFS departs from baseline expectations. This can be done either by fitting a simple demographic model or by introducing nuisance parameters that flexibly absorb demographic history and mutation-rate heterogeneity. Second, conditional on this neutral model, the SFS at selected sites (e.g., non-synonymous sites) is fitted with a parametric DFE for the population-scaled selection coefficient *N_e_s*, where *N_e_* is the effective population size and *s* is the selective disadvantage (denoted by *s_d_*) or advantage (denoted by *s_a_*) of the mutant homozygote relative to wildtype (Williamson et al. 2005; Eyre-Walker et al. 2006; Keightley and Eyre-Walker 2007; Kousathanas and Keightley 2013). Across diverse animal and plant taxa, SFS-based analyses generally support DFEs in which new mutations span a wide range of fitness effects rather than being concentrated near a single value (Gossmann et al. 2012; Galtier 2016; Chen et al. 2017; Rousselle et al. 2020).

While the DFE of nonsynonymous mutations has been investigated extensively, the contribution of mutations in non-coding regions to the DFE has received less attention. Comparative genomics across diverse taxa has revealed pervasive conservation outside exons, consistent with widespread purifying selection in non-coding DNA (Siepel et al. 2005). While only ∼7–12% of noncoding sites in mammals are conserved (Rands et al. 2014; Christmas et al. 2023; Di et al. 2025), in species that have more compact genomes, a much larger proportion of noncoding sequences may be conserved, e.g., ∼18% in *Caenorhabditis elegans*, 20-36% in *Saccharomyces cerevisiae*, and 22-55% in *Drosophila melanogaster* (Bergman and Kreitman 2001; Siepel et al. 2005; Halligan and Keightley 2006). Non-coding sequences contain cis-regulatory elements such as promoters, enhancers, silencers and insulators, untranslated regions, and diverse classes of non-coding RNAs, which collectively orchestrate when, where, and at what level genes are expressed, integrate developmental and environmental signals, and help organize higher-order chromatin architecture and gene regulatory networks (Ludwig 2002; Spitz and Furlong 2012; Siepel and Arbiza 2014; Long et al. 2016; Signor and Nuzhdin 2018; Wang et al. 2018; Keränen et al. 2022). Reflecting this central regulatory role, non-coding regulatory variants are strongly enriched among loci associated with complex traits and disease, underscoring the need to characterize the DFE of mutations in these elements (Mackay 2004; Flint and Mackay 2009; Zhang and Lupski 2015).

Although non-coding DNA plays functionally important roles, a major challenge in utilizing population genetic approaches to estimate the DFE of new mutations in these regions is the accurate identification of putatively neutral and selected sites used for inference. The identification of functionally important regulatory regions in non-coding regions is difficult, as it often requires a combination of various experimental and computational approaches (Lesurf et al. 2016; Cai et al. 2019; Keränen et al. 2022; Kosicki et al. 2025). In *Drosophila*, this challenge is compounded by the presence of short stretches (median length ∼19 bp; Bergman and Kreitman 2001) of conserved non-coding sequences and regulatory information can be densely distributed across non-coding regions ( Sabarís et al. 2024). Moreover, due to heterogeneity in the *de novo* mutation rate, recombination rate, and the effects of selection on linked sites across the genome, DFE inference methods using population genetics data are most accurate when the putatively neutral and selected sites are interdigitated (reviewed in Johri et al. 2022) so that putatively neutral sites are affected similarly to the selected sites by processes not accounted for by the methods. However, because the putatively neutral sites near functional non-coding regions are unlikely to be interdigitated with the non-coding selected sites, it is unclear whether inference will be accurate. In addition to the above, pervasive selective interference can lead to inaccurate inference of the DFE (Daigle and Johri 2025), and it is therefore not clear if interference from selected sites in nearby coding regions might affect DFE inference at regulatory regions. Finally, because regulatory regions tend to be short, few selected sites are available for inference, reducing power. Because of these challenges, we lack robust, population-genetic estimates of the DFE for regulatory non-coding mutations.

Here, we quantify the distribution of fitness effects (DFE) of regulatory non-coding DNA in natural populations of *D. melanogaster*. First, we identify putatively selected and neutral sites by collating experimentally validated regulatory regions in *D. melanogaster* and using comparative conservation data. Next, using forward simulations that incorporate realistic mutation, recombination, and genomic architecture, we ask if widely-used DFE inference methods have the power to accurately estimate the DFE of regulatory regions. While *D. melanogaster* has a highly compact genome, with potentially substantial Hill-Robertson interference (HRI), inferences were found to be minimally biased, restricted to scenarios where most mutations are mildly deleterious. We then applied this framework to three African populations to estimate the deleterious DFEs and the proportion of beneficial substitutions for coding and non-coding regions of the genome. Our results show that mutations in regulatory DNA are generally moderately deleterious, with lower—but still substantial—fractions of beneficial substitutions compared to coding regions. Incorporating these empirically grounded non-coding DFEs into maps of background selection, we demonstrate improved predictions of genome-wide diversity, highlighting the importance of accurately modeling selection across the genome.

## RESULTS

### Population stratification and inversion filtering in African *D. melanogaster* populations

To investigate the distribution of fitness effects (DFE) in non-coding regions, we utilized *D. melanogaster* single nucleotide polymorphism (SNP) data from chromosomes 2 and 3 provided by Coughlan *et al*. (2022). Due to the underlying assumptions of panmixia made by most DFE inference methods, we attempted to identify populations that have minimal population substructure and the least amount of admixture. We therefore specifically focused on populations from sub-Saharan Africa–the presumed ancestral range of the species (Pool et al. 2012; Lack et al. 2015; Mansourian et al. 2018; Sprengelmeyer et al. 2020), as these individuals formed distinct, monophyletic clusters with minimal signs of admixture compared to other populations (see Figure 1 from Coughlan *et al*. (2022)). Focusing on ancestry groups identified using *PCAngsd* in Coughlan *et al*. (2022), we restricted our analyses to individuals whose geographic sampling locations matched their inferred ancestry clusters. *PCAngsd* performs principal component analysis directly on genotype likelihoods rather than the genotype calls to capture major axes of genetic variation and then clusters individuals in that reduced PC space to infer ancestry groups (Meisner and Albrechtsen 2018). These populations were defined as “South,” consisting entirely of individuals sampled in or near Mana Pools National Park in Zimbabwe; “East,” comprising individuals from Rwanda and Uganda; and “West,” including individuals from Cameroon, Gabon, Ghana, Guinea, and Nigeria.

**Figure 1:**
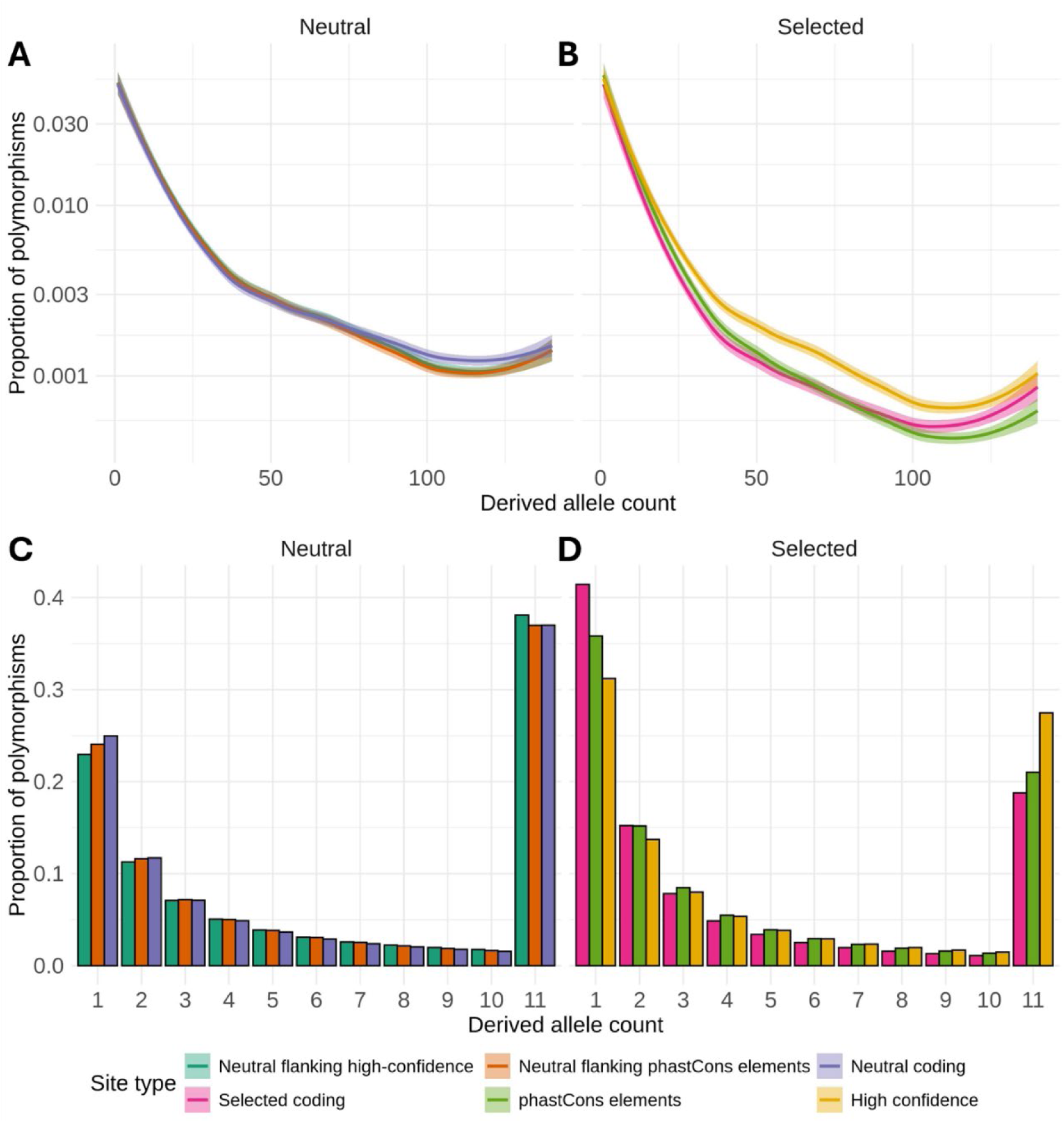
Empirical unfolded site frequency spectra of putatively neutral and selected alleles from 144 individuals from the South population. The *y*-axis represents the proportion of segregating polymorphisms that fall into the given derived allele count on the *x*-axis. “Neutral flanking” sites refer to putatively neutral sites in non-coding regions lie within 5 kb of the corresponding selected sites and do not include 4-fold degenerate sites. For the top panels the y-axis is plotted on a log scale, with all derived allele counts displayed. Lines were smoothed with LOESS. For the bottom panels the last class (11) refers to the derived allele counts 11 and above.

We assessed differentiation between and within these populations by calculating pairwise *F_ST_* values using fourfold degenerate sites from coding regions. Flies in the East population showed no evidence of differentiation between individuals sampled from Uganda and Rwanda (*F_ST_* =-0.018, 95% CI: -0.0240 to -0.0128, *p*=1.0). In the West population, most *F_ST_* values were near zero or negative, indicating little or no differentiation; however, individuals from Ghana exhibited mild differentiation (*F_ST_*=-0.024 to -0.0128, *p* < 10^−14^, Table S1) from other West individuals, prompting us to exclude them from subsequent analyses. In total, we retained 141 individuals in South, 29 in East, and 26 in West, sequenced with short reads at average depths ranging from 15× to 145×. Differentiation among the three focal populations was significant in all pairwise comparisons, with South–West showing the highest *F_ST_* (0.0975, 95% CI: 0.0833–0.1116), followed by South–East (0.0781, 95% CI: 0.0630–0.0931) and East–West (0.0391, 95% CI: 0.0327–0.0456; all *p* < 10^−23^).. These patterns are consistent with the geographic distance between the populations and previous analyses of differentiation(using all SNPs) among individuals sampled from these regions (Coughlan et al. 2022; J. Chen et al. 2024).

Large polymorphic inversions have been previously implicated in driving differentiation within *Drosophila* populations (Krimbas and Powell 1992; Corbett-Detig and Hartl 2012). As major inversions were genotyped by Coughlan *et al*. (2022), we calculated pairwise *F_ST_* values between homozygous inversion-present and homozygous inversion-absent individuals within each subpopulation, comparing all inversions for which the homozygous inversion genotype was observed in more than one individual in that subpopulation. It is important to note that regions within and flanking major inversions were filtered out of the Coughlan *et al*. (2022) dataset, removing the regions most likely differentiated due to inversions. In the East population, the inversion *ln(3R)P_e* was at a frequency of 17%, showing no differentiation between individuals with and without the inversion (*F_ST_* = −0.0188, 95% CI: -0.0230 to -0.0145). In the West population, we also found almost no differentiation between individuals with and without the inversions *In(2L)t* (*F_ST_* = 0.0041, 95% CI: −0.0052 to 0.0109), *In(2R)NS* (*F_ST_* = −0.0718, 95% CI: −0.0887 to −0.0724), *In(3L)P* (*F_ST_* = −0.0236, 95% CI: −0.0326 to −0.0147), *In(3R)k* (*F_ST_* = −0.0033, 95% CI: −0.0067 to 0.0001), and *In(3R)P_e* (*F_ST_* = −0.015, 95% CI: −0.0183 to −0.0109). In the South population, *In(3L)OK* is nearly fixed, with only one homozygous absent individual in our sample, so it did not meet our threshold for calculating *F_ST_*. Taken together, these analyses suggest that our final dataset is unlikely to be strongly influenced by admixture, inversions, or extensive population structure.

The South population exhibited higher mean nucleotide diversity (*π* = 0.016) and Watterson’s theta (*θ*_W_ = 0.020) at 4-fold degenerate sites compared to the East (*π* = 0.011; *θ*_W_ = 0.012) and West (*π* = 0.011; *θ*_W_ = 0.012) populations, suggesting a larger effective population size (Table 1). In addition, mean Tajima’s *D* values were more negative in the south (-0.644) than in East and West (-0.452 and -0.477, respectively), indicating an excess of rare variants that may reflect stronger recent population growth and/or stronger effects of selectionat linked sites. In contrast, the ratio of nucleotide diversity at 0-fold degenerate over 4-fold degenerate sites 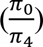 was similar across all three populations—with a slight elevation in the West population (0.141 vs. 0.134 for South and 0.135 for East)—implying that the overall strength of purifying selection on amino-acid changes was relatively consistent.

**Table 1:**
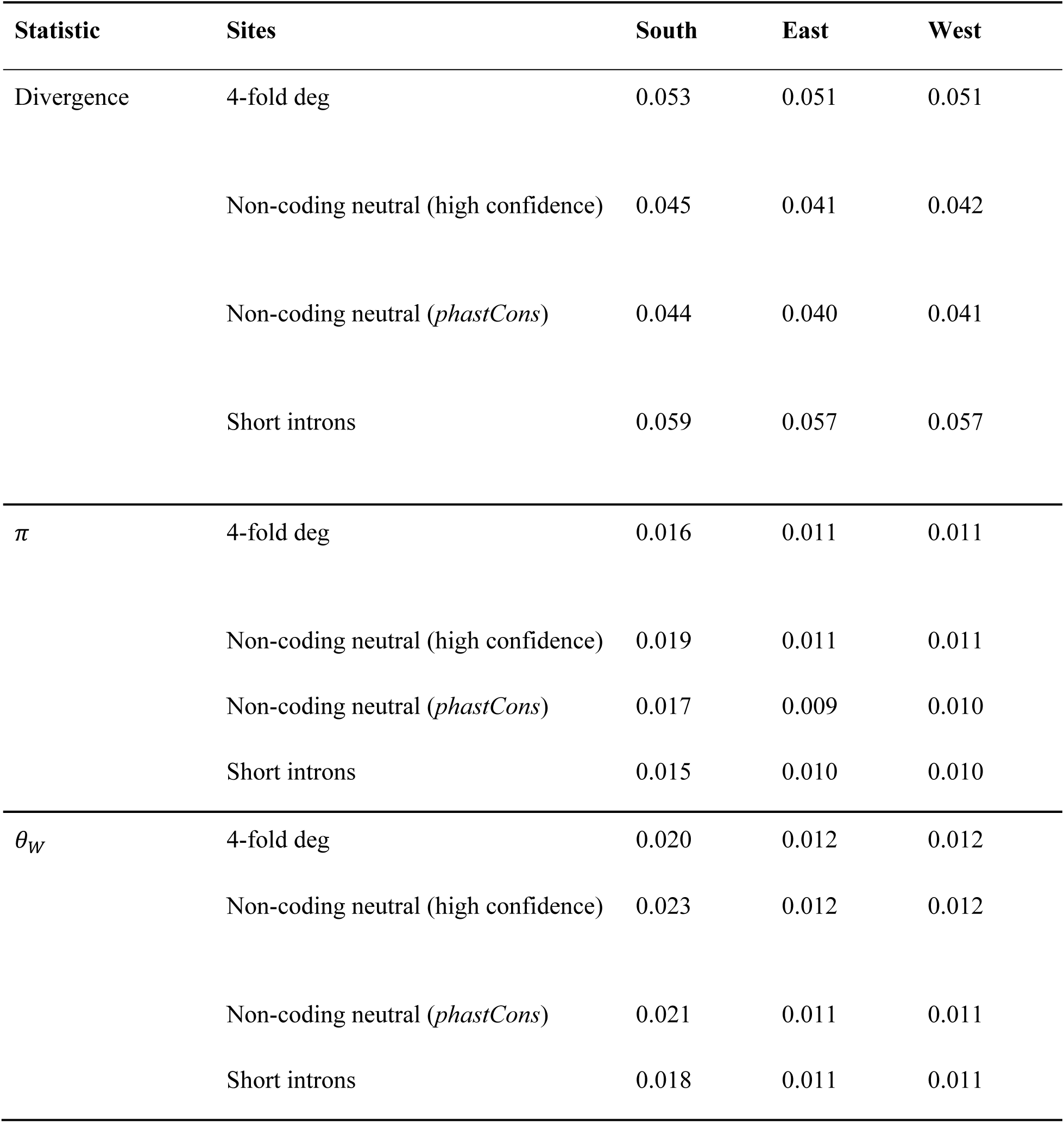
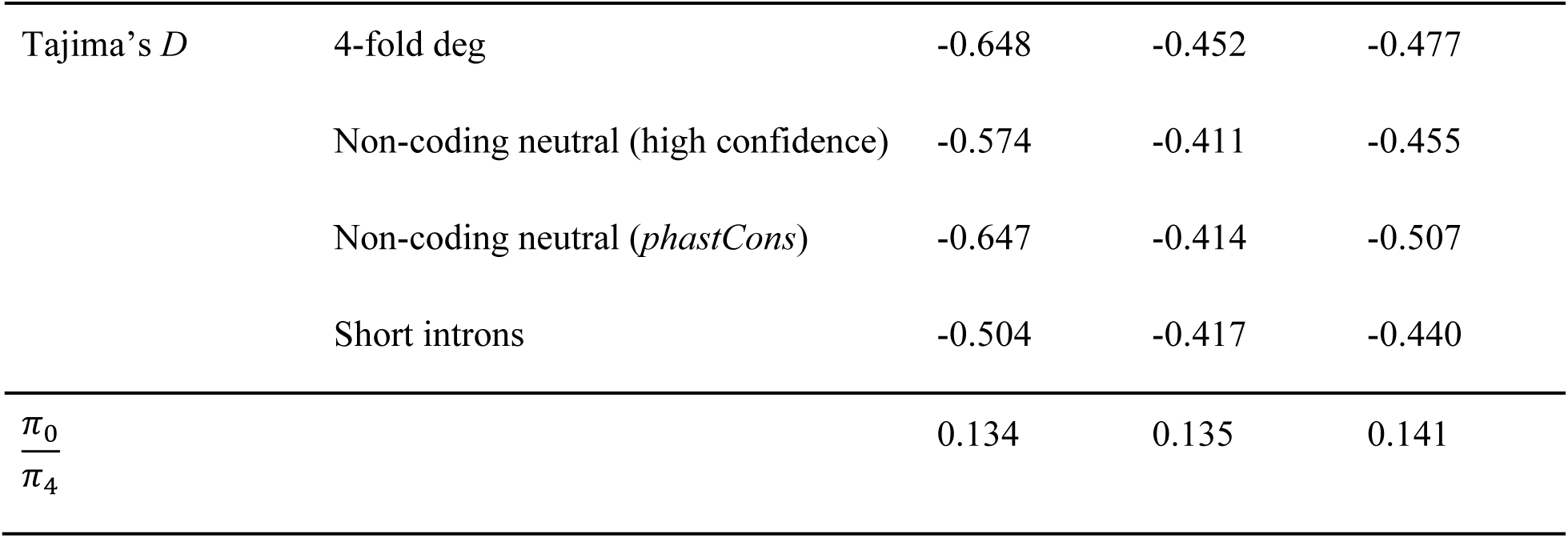
Summary statistics describing the three populations used for DFE inference. Non-coding neutral (high confidence) and non-coding neutral (*phastCons*) refers to neutral sites in non-coding regions adjacent (within 5 kb on either side) to high confidence regions annotated in this study and *phastCons* elements, respectively.

| Statistic | Sites | South | East | West |
| --- | --- | --- | --- | --- |
| Divergence | 4-fold deg | 0.053 | 0.051 | 0.051 |
|  | Non-coding neutral (high confidence) | 0.045 | 0.041 | 0.042 |
|  | Non-coding neutral (*phastCons*) | 0.044 | 0.040 | 0.041 |
|  | Short introns | 0.059 | 0.057 | 0.057 |
| $\pi$ | 4-fold deg | 0.016 | 0.011 | 0.011 |
|  | Non-coding neutral (high confidence) | 0.019 | 0.011 | 0.011 |
|  | Non-coding neutral (*phastCons*) | 0.017 | 0.009 | 0.010 |
|  | Short introns | 0.015 | 0.010 | 0.010 |
| $\theta_w$ | 4-fold deg | 0.020 | 0.012 | 0.012 |
|  | Non-coding neutral (high confidence) | 0.023 | 0.012 | 0.012 |
|  | Non-coding neutral (*phastCons*) | 0.021 | 0.011 | 0.011 |
|  | Short introns | 0.018 | 0.011 | 0.011 |
| Tajima's $D$ | 4-fold deg | -0.648 | -0.452 | -0.477 |
|  | Non-coding neutral (high confidence) | -0.574 | -0.411 | -0.455 |
|  | Non-coding neutral (*phastCons*) | -0.647 | -0.414 | -0.507 |
|  | Short introns | -0.504 | -0.417 | -0.440 |
| $\frac{\pi_0}{\pi_4}$ | | 0.134 | 0.135 | 0.141 |

### Identifying selected and putatively neutral sites in non-coding genomic regions

To infer the DFE in non-coding regulatory regions, it is important to identify (1) non-coding sites that are functionally important, and (2) putatively neutral sites that can be adequately used for inference. To identify functional non-coding regions, we employed the *REDfly* database—a curated collection of experimentally validated transcriptional cis-regulatory modules (CRMs) and transcription factor binding sites (TFBSs) in *Drosophila melanogaster* (Keränen et al. 2022). Rather than depending on a single methodology, *REDfly* combines results from multiple complementary experimental approaches, including chromatin immunoprecipitation followed by sequencing (ChIP-seq) assays, *DNase I* hypersensitivity mapping, electrophoretic mobility shift assays, enhancer fluorescence activated cell sorting (FACS) sequencing, and in vivo reporter assays. We refer to these experimentally validated regions as “high-confidence” regions, which represented 0.2% of the genome and include enhancers, promoters, TFBS, and miRNAs. This small fraction reflects the limited genomic coverage of experimentally validated annotations in REDfly.

As these high-confidence regions represent such a small fraction of the genome, we included two additional categories of potentially functional non-coding sites. First, we included *phastCons* elements (Siepel et al. 2005), which are conserved sequences identified from alignments of 27 insect species related to *D. melanogaster*. Because conservation-based annotations preferentially identify sequences subject to long-term purifying selection, they may exclude lineage-specific or rapidly evolving regulatory elements. Second, we included computationally predicted and inferred CRMs from the *REDfly* database and additional computational annotations from *FlyBase* and *ORegAnno*, referring to these putatively functional regions as “low-confidence” regions. In our final annotation, coding (17.1% of the genome), untranslated regions (UTRs; 8% of the genome), high-confidence (0.2%), *phastCons* elements (35.5%), and low-confidence (26.2%) regions comprised non-overlapping sites and totaled 87.3% of the genome (see *Methods*). As we were confident that UTRs, high-confidence regions, and *phastCons* elements represented non-coding sites under direct selection, DFE inference was restricted to those regions. In order to identify putatively neutral non-coding sites across the genome, we excluded all sites that could potentially be under direct selection, i.e., the high- and low-confidence sites, coding regions, UTRs, and *phastCons* elements. The remaining unannotated genomic sites, which comprised the remaining 13% of the genome, were considered potentially neutral intergenic sites. To further refine this neutral set, we applied single-site *phastCons* scores from the same alignment, which assess conservation at individual nucleotides; sites with scores above 0.1 were excluded to ensure neutrality (see Methods for a comparison of various cutoffs). This filtering resulted in our final set of putatively neutral sites in non-coding regions that covered 3.2% of the genome. Given that DFE inference approaches assume similar mutation and recombination rates between selected and neutral sites, and perform the best when selected and neutral sites are interdigitated (Johri et al. 2022), we further restricted our neutral sites to those in the 5kb flanking regions up- and downstream of our high-confidence and *phastCons* regions, giving us a final set of putatively neutral sites for inference (see Table S2). Nucleotide diversity (*π*) at the final set of putatively neutral non-coding sites in the South population was found to be 0.017-0.019, similar to levels at 4-fold degenerate sites, 0.016, suggesting that these sites are likely not under selection. Divergence from *D. simulans* at neutral non-coding sites was found to be ∼0.044 in all populations, which was slightly lower than the value at synonymous sites (Table 2). This could indicate weak selection acting on the non-coding neutral sites.

**Table 2:** Estimates of the distribution of fitness effects of new deleterious mutations (reported as the mean of population-scaled selection coefficients, 2*N_e_s̅_d_* and the proportion of beneficial substitutions (*α*) in coding and conserved non-coding regions across a range of species with available estimates. Here *N_e_* is the effective population size and *s̅_d_* is the mean selective disadvantage of the mutant homozygote.

| Species | Coding<br>$2N_e\overline{s_d}$ | Coding<br>$\alpha$ | Conserved non-coding $2N_e\overline{s_d}$ | Conserved<br>non-coding $\alpha$ | Reference |
| --- | --- | --- | --- | --- | --- |
| *Drosophila melanogaster* | 555-2294 | 0.47-0.557 | ~50 | ~0.30 | This study |
| *Drosophila melanogaster* | - | - | 20-200 | 0.15-0.19 | Andolfatto 2005;<br>Casillas et al. 2007 |
| *Mus musculus castaneus* | 2000 | 0.3 | 80-90 | 0.15-0.19 | Halligan et al. 2013;<br>Booker and<br>Keightley 2018;<br>Booker et al. 2021 |
| *Homo sapiens* | 328 | - | 16-27 | 0.0467 | Torgerson et al.<br>2009; Kim et al.<br>2017; Di et al. 2025 |
| *Capsella grandiflora* | 404 | 0.41 | 31 | 0.54 | Williamson et al.<br>2014 |

To further confirm if our set of identified neutral non-coding sites might experience weak selection, we obtained unfolded site frequency spectra for each population, inferring ancestral states at both polymorphic and invariant sites using *D. simulans* as an outgroup (Sendrowski and Bataillon 2024). Encouragingly, the spectra for our synonymous and non-coding neutral sites were highly similar for low- and high-frequency alleles, suggesting that our non-coding neutral sites are minimally affected by selection, i.e., no more than 4-fold degenerate sites (Figure 1). Short introns, which are commonly used as a neutral reference in *D. melanogaster*, showed similarly shaped SFS and comparable nucleotide diversity and Tajima’s *D*. However, divergence in short introns was somewhat higher than that of our non-coding neutral sites (Table 1), suggesting that the set of non-coding neutral sites might include some selected sites or may be biased towards mutational coldspots. All putatively selected coding and non-coding regions had more rare alleles in the SFS relative to neutral sites, consistent with direct selection, with coding regions appearing to be under stronger purifying selection than non-coding regions (Figure 1). Reflecting these results, nucleotide diversity (*π*), Watterson’s theta (*θ*_W_), and Tajima’s *D* were similar within all populations for putatively neutral sites in coding and non-coding regions (Table 1).

### Testing the accuracy and power of DFE-inference in non-coding regions

We tested the performance of three commonly used DFE-inference programs—*DFE-alpha* (Keightley and Eyre-Walker 2007), *GRAPES* (Galtier 2016), and *fastDFE* (Sendrowski and Bataillon 2024) using realistic forward-in-time simulations of *D. melanogaster* -like populations. These methods differ slightly in their modeling assumptions and treatment of deleterious and beneficial mutations. Our simulations incorporated empirically derived recombination maps, variation in mutation rate, and the exact positions of genomic elements under selection (Figure 2). We modeled five annotation classes: coding exons, high-confidence regulatory regions, *phastCons* elements, low-confidence regulatory regions, and neutral (details in Table S2), where the first three classes represent directly selected sites. For mutations arising at directly selected sites, we assumed that 99.98% were deleterious and the remaining 0.02% were beneficial. While the DFE at coding and low-confidence regulatory regions was fixed (see Methods), the deleterious DFE assigned to high-confidence regulatory sites was varied from predominantly weakly, moderately, or strongly deleterious mutations and was the one inferred. Beneficial effects were drawn from an exponential distribution with mean 2*N_e_s̅_a_* = 250 following Campos et al. (2017), who estimated the rate and strength of favorable mutations in *D. melanogaster* using a model incorporating background selection and selective sweeps that included the effects of gene conversion. Because accurately estimating the strength and proportion of beneficial mutations is difficult, but not their product (which is proportional to *α*), for the same value of *α*, we simulated two additional DFEs where beneficial mutations were weaker and more common (2*N_e_s̅_a_* = 25; *fpos* = 0.2%) or stronger and rarer (2*N_e_s̅_a_* = 2500; *fpos* = 0.002%).

**Figure 2:**
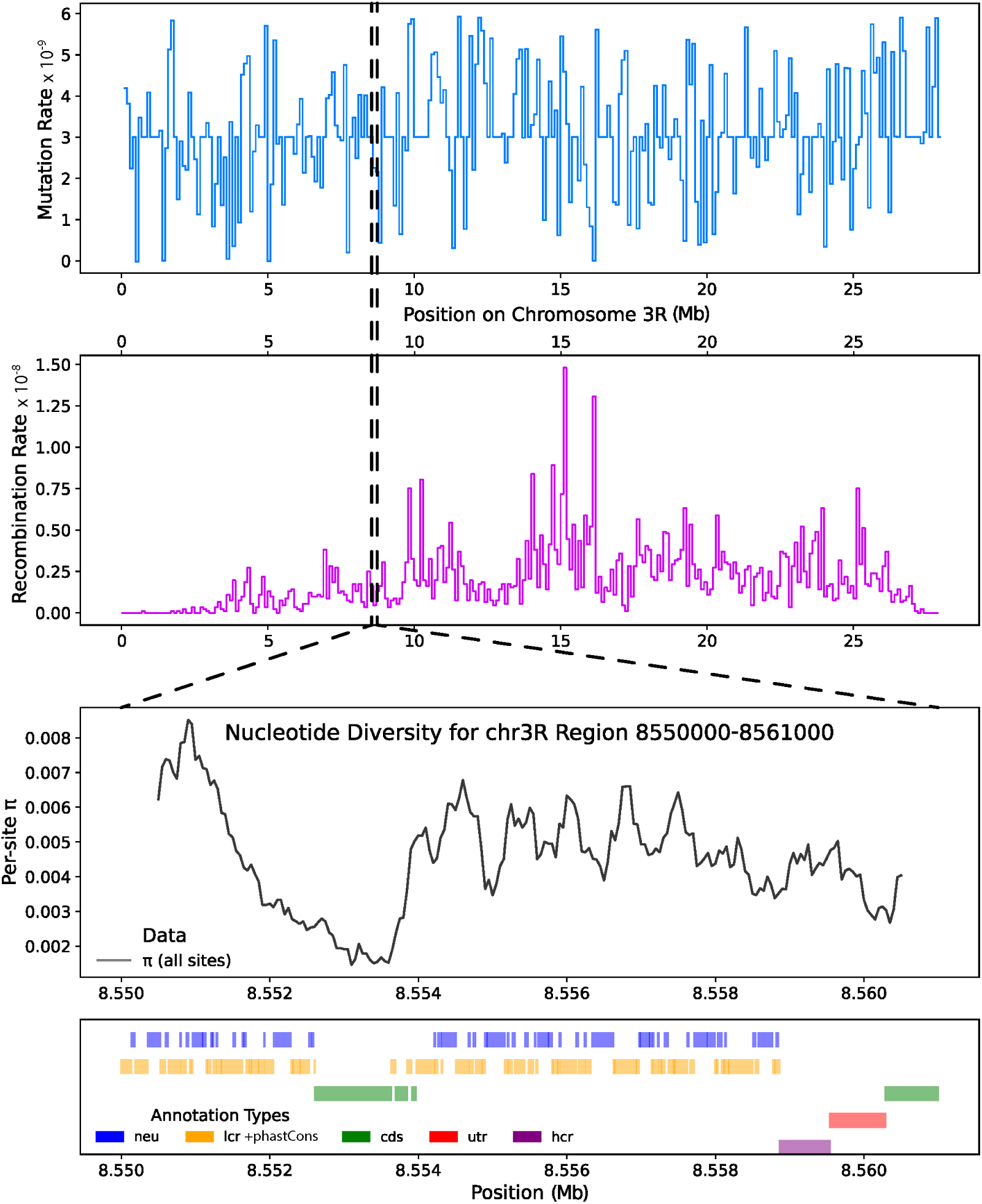
A visualization of the framework used for forward-in-time simulations. Top panels show the mutation-rate (top) and crossing-over-rate (middle) landscapes used in the forward simulations along chromosome arm 3R, parameterized from empirical *D. melanogaster* maps and applied piecewise across 100kb windows. Dashed vertical lines mark the focal region shown below. The lower panel shows a zoomed-in segment (chr3R: 8,550,000–8,561,000) illustrating the simulated annotation architecture and the resulting per-site nucleotide diversity (π) from the simulation output. Annotation tracks indicate the positions of neutral sites (neu; blue), *phastCons* + low-confidence regulatory regions (lcr) (annotated together in orange), coding sequence (cds; green), UTRs (utr; red), and high-confidence regulatory regions (hcr; purple).

#### Accuracy of inference of the deleterious DFE

We tested the inference of the DFE using the folded SFS, as this mitigates potential issues with incorrectly polarized polymorphisms in our empirical data (Tataru et al. 2017; Keightley and Jackson 2018). In general, all methods improved in performance as the number of sites increased, with the standard deviation of the inferred proportion of DFE classes decreasing (Figure 3), with more than 10,000 selected sites (and 7,193 flanking neutral sites), required for reliable DFE inference, with or without beneficial mutations (Figure 3; Figure S1). Note that we discuss the accuracy of inference with respect to the number of selected *sites*, not the number of *variants* because this quantity is fixed by the simulated genomic architecture, whereas the number of segregating selected variants is an outcome of the simulation and varies substantially across DFEs and sample sizes. The DFEs comprising primarily moderately and strongly deleterious mutations were estimated well by all methods, while the DFE comprising mostly weakly deleterious mutations was more difficult to estimate, with *fastDFE* and *GRAPES* having more accurate estimates than *DFE-alpha*. In particular, the proportion of weakly deleterious mutations was underestimated and mutations were redistributed into the effectively neutral and moderately deleterious classes. This bias increased as beneficial mutations became stronger and rarer, while holding *f_pos_* × 2*N_e_s̅_a_* constant (Figure S2), suggesting that the effects of selection on linked sites are likely responsible for the observed inaccuracy.

**Figure 3:**
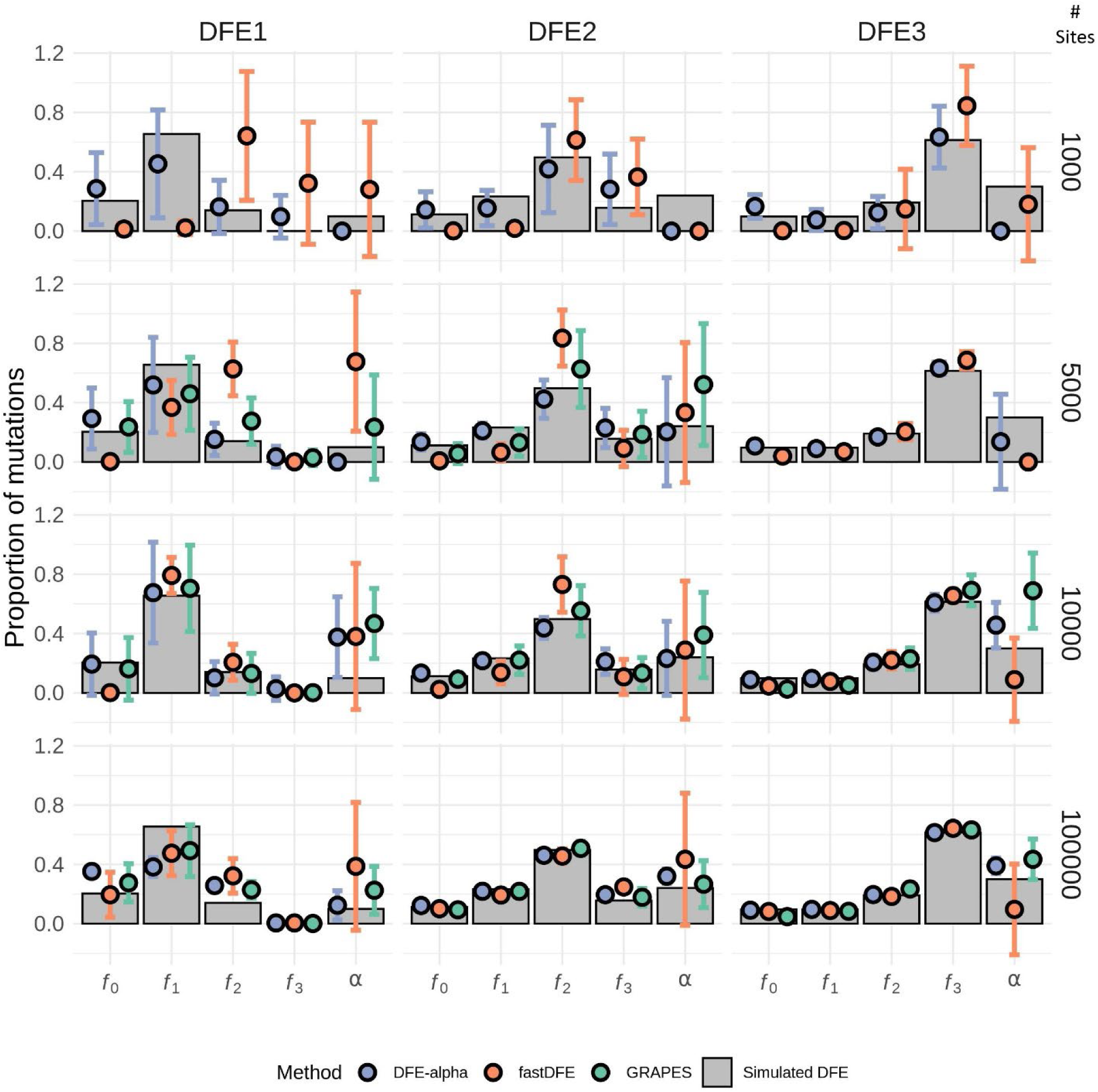
Performance of *DFE-alpha*, *fastDFE*, and *GRAPES* in estimating the DFE of simulated high-confidence regulatory regions for varying numbers of selected sites and DFEs. All inferences were performed with 200 diploid individuals using the folded SFS as input. Comparisons of the simulated and inferred DFE are shown in terms of the proportion of mutations, *f*_0_, *f*_1_, *f*_2_, and *f*_3_, in effectively neutral (0 ≤ 2*N_e_s_d_* < 1), weakly deleterious (1 ≤ 2*N_e_s_d_* < 10), moderately deleterious (10 ≤ 2*N_e_s_d_* < 100), and strongly deleterious (100 ≤ 2*N_e_s_d_*) classes of mutations, respectively, along with the proportion of substitutions that are beneficial (*α*). The simulated DFE is denoted with grey bars, and the inferred DFE is shown as circles. In all panels, error bars denote the standard deviation of estimates across 10 independent subsampling replicates, in which sites were randomly drawn without replacement from the full set of simulated sites. The number of selected sites included in each replicate is indicated on the right-hand *y*-axis.

Generally, the inferred mean 2*N_e_s̅_d_* and shape (*β*) of the deleterious DFE did not strongly depend on the number of individuals used for inference (Figures S3 and S4), except when inferring the DFE comprising primarily weakly deleterious mutations (where estimates of *β* by *GRAPES* and *fastDFE* improved with more individuals). While all methods approximated the shape of the DFE well across different numbers of individuals (ranging from 25-200) and 10,000 selected sites were sufficient to provide a reasonably accurate inference, the DFE with mostly weakly deleterious mutations remained the most challenging to estimate. Overall, *DFE-alpha* provided the most consistent estimates of the deleterious DFE across scenarios, with the caveat that it was less accurate for the DFE skewed towards weakly deleterious mutations.

#### Accuracy of inference of the proportion of beneficial substitutions

(*α*): Estimates of the proportion of beneficial substitutions (*α*) were calculated differently by the three methods, with *DFE-alpha* and *GRAPES* utilizing divergence data and *fastDFE* relying on its fitted distribution of fitness effects alone (see Methods). With 100,000 selected sites, *α* was estimated accurately by both *DFE-alpha* and *GRAPES* in most scenarios, while *fastDFE* produced inaccurate predictions with high variance across replicates (Figure 3). However, in simulations with stronger and rarer beneficial mutations, *GRAPES* greatly overestimated *α*, unlike *DFE-alpha* (Figure S2). With 10,000 selected sites, *α* tended to be overestimated by all programs, though *DFE-alpha* tended to produce lower estimates than *GRAPES*. A similar pattern was observed when varying the number of individuals, though *GRAPES* performed particularly poorly with lower numbers (<=50) of individuals for the DFE skewed towards moderately deleterious mutations. In the absence of beneficial mutations *GRAPES* estimated a non-zero positive mean value of *α* in 44% of cases (Figure S1). *FastDFE*, on the other hand, consistently overestimated *α*, with the majority of its estimates exceeding 0.25. In summary, *DFE-alpha* provided the most reliable *α* estimates across all scenarios tested, especially when 100,000 selected sites were used. When the unfolded SFS was used as input instead of the folded SFS, the parameters of the deleterious and beneficial DFE were inferred with similar accuracy for *GRAPES* and *fastDFE*, while *DFE-alpha* predictions were worse due to the overestimation of *γ* and the underestimation of *β*, as expected because *DFE-alpha* does not account for segregating beneficial alleles in the SFS by default (Figure S5).

#### Considering changes in outgroup population size

The model-based DFE inference methods used here estimate *α* by comparing the observed nonsynonymous divergence to the expected nonsynonymous divergence under a model containing only neutral and deleterious mutations. Because divergence is often estimated as pairwise differences between two taxa (the focal and outgroup species), the expected number of neutral and weakly deleterious fixations is highly dependent on the assumed ancestral and outgroup population sizes of both taxa. As such, misspecification of outgroup population sizes can bias *α* estimates by altering the expected number of weakly deleterious substitutions on the outgroup lineages (Zhen et al. 2021). For instance, if the outgroup population is assumed to be smaller than the actual size, more fixations at non-synonymous sites would be attributed to weakly deleterious mutations than otherwise, and therefore the true proportion of beneficial fixations would be underestimated. In general, assuming an outgroup population size that is too small biases *α* downward, whereas assuming sizes that are too large can bias *α* upward. Note that such biases should ideally be present only when divergence is estimated pairwise using two species, not when correctly assigned branch-specific fixations are used to calculate divergence. To test this with parameters relevant to the *D. melanogaster–D. simulans* demography, we simulated a split with constant population sizes and an outgroup ∼50% larger than the ingroup (following Andolfatto et al. 2011). Because the assumed value of the ancestral population size also substantially determines estimates of *α*, we varied the ancestral population size across three scenarios (0.5, 1, and 2 million individuals). As expected, the true proportion of beneficial fixations differed between the branches leading to the two taxa. Across the three scenarios, *α*_mel_ was equal to 0.462, 0.440, and 0.447 and *α*_sim_ was equal to 0.574, 0.568, and 0.562, respectively. The corresponding true pairwise *α* values, calculated as the proportion of beneficial substitutions among all substitutions across both lineages, were 0.523, 0.509, and 0.509. *DFE-alpha* estimates were 0.498 ± 0.035, 0.523 ± 0.023, and 0.534 ± 0.043, respectively, closely approximating the pairwise values but overestimating lineage-specific *α* for *D. melanogaster*. *GRAPES* produced higher and more variable estimates of 0.674 ± 0.195, 0.632 ± 0.202, and 0.626 ± 0.231 (mean ± SD across ten simulation replicates per scenario; Figure S6). Therefore, our simulations suggest that estimates of lineage-specific α in *D. melanogaster* using *DFE-alpha* may be overestimated by a mean of 15.4%, depending on the demographic scenario.

### Inferring the DFE of non-synonymous sites in *D. melanogaster*

We inferred the DFE for non-synonymous sites using the folded SFS in each *D. melanogaster* population using *DFE-alpha*, *fastDFE*, and *GRAPES* The mean population-scaled selection coefficient (2*N_e_s̅_d_*) of the gamma distribution estimated by each method for each population varied considerably (between -555 and -2294), while the inferred *β* of the gamma distribution was more consistent (0.304-0.355). Across these estimates, SDs among the 10 subsampling replicates generally remained small, ranging from 27.0—246 for 2*N_e_s̅_d_* and 0.005—0.021 for *β* (Table S4). Because these fitted gamma parameters are not independent, they can vary even when the proportion of weakly vs strongly deleterious mutations remains similar across populations/replicates (Chen et al. 2017; Galtier and Rousselle 2020). We therefore summarize the inferred DFE as the proportion of mutations in four population-scaled deleterious-effect classes: effectively neutral (0 ≤ 2*N_e_ s_d_* < 1), weakly deleterious (1 ≤ 2*N_e_s_d_* < 10), moderately deleterious (10 ≤ 2*N_e_ s_d_* < 100), and strongly deleterious (100 ≤ 2*N_e_s_d_*), where *s_d_* refers to the selective disadvantage of the mutant homozygote. Across methods and populations, a large proportion (> 60%) of mutations were estimated to be strongly deleterious by all methods (Figure 4; Figure S7). Our inference of a strongly deleterious coding DFE in the new African *D. melanogaster* samples is broadly consistent with previous estimates from African populations using similar methods (Keightley and Eyre-Walker 2007; Shapiro et al. 2007; Kousathanas and Keightley 2013; Huber et al. 2017; Ragsdale and Gutenkunst 2017), including studies of individuals from Zambia (Keightley and Eyre-Walker 2007; Shapiro et al. 2007; Huber et al. 2017; Ragsdale and Gutenkunst 2017), Zimbabwe (Keightley and Eyre-Walker 2007; Shapiro et al. 2007), and Rwanda (Kousathanas and Keightley 2013), all of which estimated a strongly deleterious DFE (2*N_e_s̅_d_* and *β*∼0.3 − 0.38). Estimates of the proportion of beneficial fixations (*α*) in coding regions ranged from 0.47 (*DFE-alpha*, West population) to 0.557 (*GRAPES*, South population), with estimates for all three populations falling close to 0.5, which replicates previous estimates of this proportion in African *D. melanogaster* populations using similar methods (Elyashiv et al. 2016; Keightley et al. 2016; Lange and Pool 2018; Zhen et al. 2021; Marsh and Johri 2024). However, note that because the outgroup population is larger in size than assumed by the employed inference methods, estimates of *α* could be ∼0.4.

**Figure 4:**
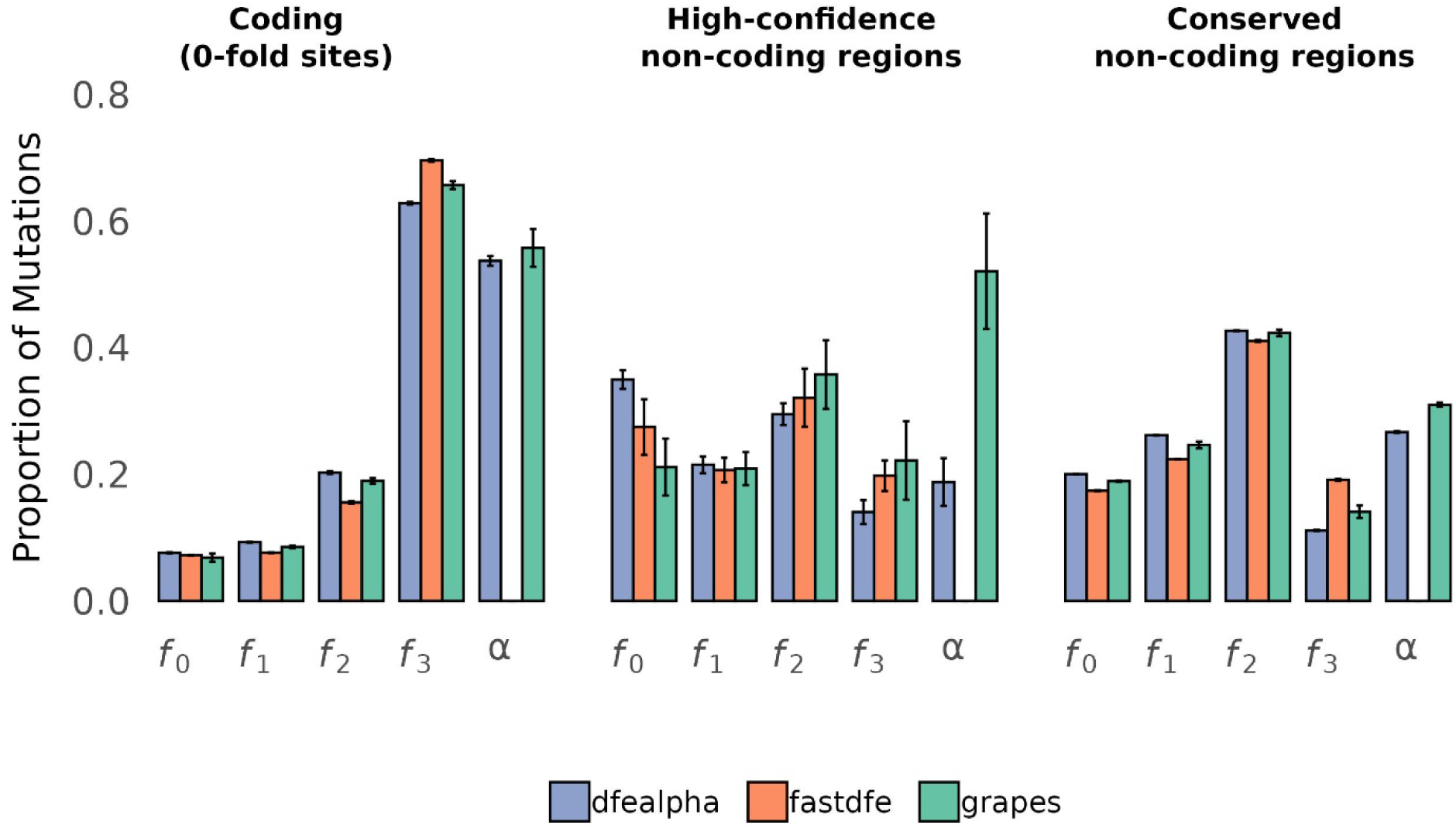
Inference of the distribution of fitness effects (DFE) of new mutations in the South population of *D. melanogaster* using the folded SFS as input for coding, high-confidence regulatory regions, and conserved non-coding regions. The DFE is shown in terms of the proportion of mutations (*f*_0_, *f*_1_, *f*_2_, *f*_3_) in effectively neutral (0 ≤ 2*N_e_s_d_* < 1), weakly deleterious (1 ≤ 2*N_e_s_d_* < 10), moderately deleterious (10 ≤ 2*N_e_s_d_* < 100), and strongly deleterious (100 ≤ 2*N_e_s_d_*) classes of mutations, along with the proportion of substitutions that are beneficial (*α*). *α* predictions from *fastDFE* were excluded due to their inaccuracy in the benchmarking simulations.

### Inferring the DFE of non-coding regulatory regions in *D. melanogaster*

We performed inference in non-coding regulatory elements that had more than 10,000 selected sites when inferring the deleterious DFE and used all individuals (27-141) from each population and did not employ *fastDFE* when estimating the proportion of new beneficial substitutions due to its poor performance on simulated data. The inferred DFEs for high-confidence regions were much less deleterious than for non-synonymous sites, with high-confidence regions showing an approximately even distribution across the four deleterious-effect classes (∼25% in each class), in contrast to non-synonymous sites, where ∼65% of new mutations were estimated to be strongly deleterious. The exact parameters of the inferred gamma distribution varied by the method and population (with 2*N_e_s̅_d_* varying from -42 to -99 and *β* from 0.21 to 0.32, SDs between 14.8-59.7 and 0.019-0.257), but there was no consistent difference in DFEs across populations (Figure S7; Figure S8). *DFE-alpha* estimated the proportion of beneficial substitutions as 0.34, 0.26, and 0.19 for the East, West, and South populations, respectively, while *GRAPES* produced estimates of 0.30, 0.37, and 0.52, respectively. Because these populations split relatively recently, they are expected to share most fixations. Thus differences in the estimated proportion of beneficial substitutions are likely due to uncertainty in the inference.

We additionally inferred the DFE for conserved non-coding regions as defined by *phastCons* elements, which lack experimental validation for function, but make up a much larger proportion of the genome than the high-confidence regions. The DFE for conserved non-coding regions had a larger proportion of mutations in the moderately deleterious class compared to the DFE in high-confidence regions (Figure 4). Overall, the inferred DFEs for non-coding regions were highly similar across populations. The largest difference was observed for conserved non-coding regions, where the West population showed a slight shift from the strongly deleterious class toward the weakly and moderately deleterious classes relative to the East and South populations (Figure S7).

As a sanity check, we inferred the DFE for the low-confidence non-coding regions (Figure S9). Their DFE was found to be substantially skewed towards neutral mutations, with approximately 50% of mutations inferred to be effectively neutral across methods. However, these methods also inferred a relatively large strongly deleterious class (28.4% by *DFE-alpha* in the South population), compared with 14.2% in high-confidence regions, producing a more bimodal DFE. Because our simulations showed reduced accuracy when the true DFE was dominated by effectively neutral and weakly deleterious mutations, we interpret these estimates cautiously.

To compare the selective effects of mutations in different types of regulatory regions, we subdivided the high-confidence regions from the *REDfly* database into transcription factor binding sites, promoters, and enhancers, inferring DFEs only in the South population. The deleterious DFEs for promoters (Figure 5A) and enhancers (Figure 5B) resembled those for all combined high-confidence regions, with ∼25% of mutations distributed across the four DFE classes, though they produced more variable results across subsampling replicates compared to other regions, likely due to limited flanking neutral sites (∼9k; Table S2). The deleterious DFE inferred for UTR regions was centered around moderately deleterious mutations, similar to what we observed in high-confidence and conserved non-coding regions, but with higher 2*N_e_s̅_d_* values (Figure 5C). Notably, the *REDfly* transcription factor binding sites showed a more strongly deleterious DFE compared to other high-confidence regions (2*N_e_s̅_d_* between -308 (*GRAPES*) and -1165 (*fastDFE*); Figure 5D).

**Figure 5:**
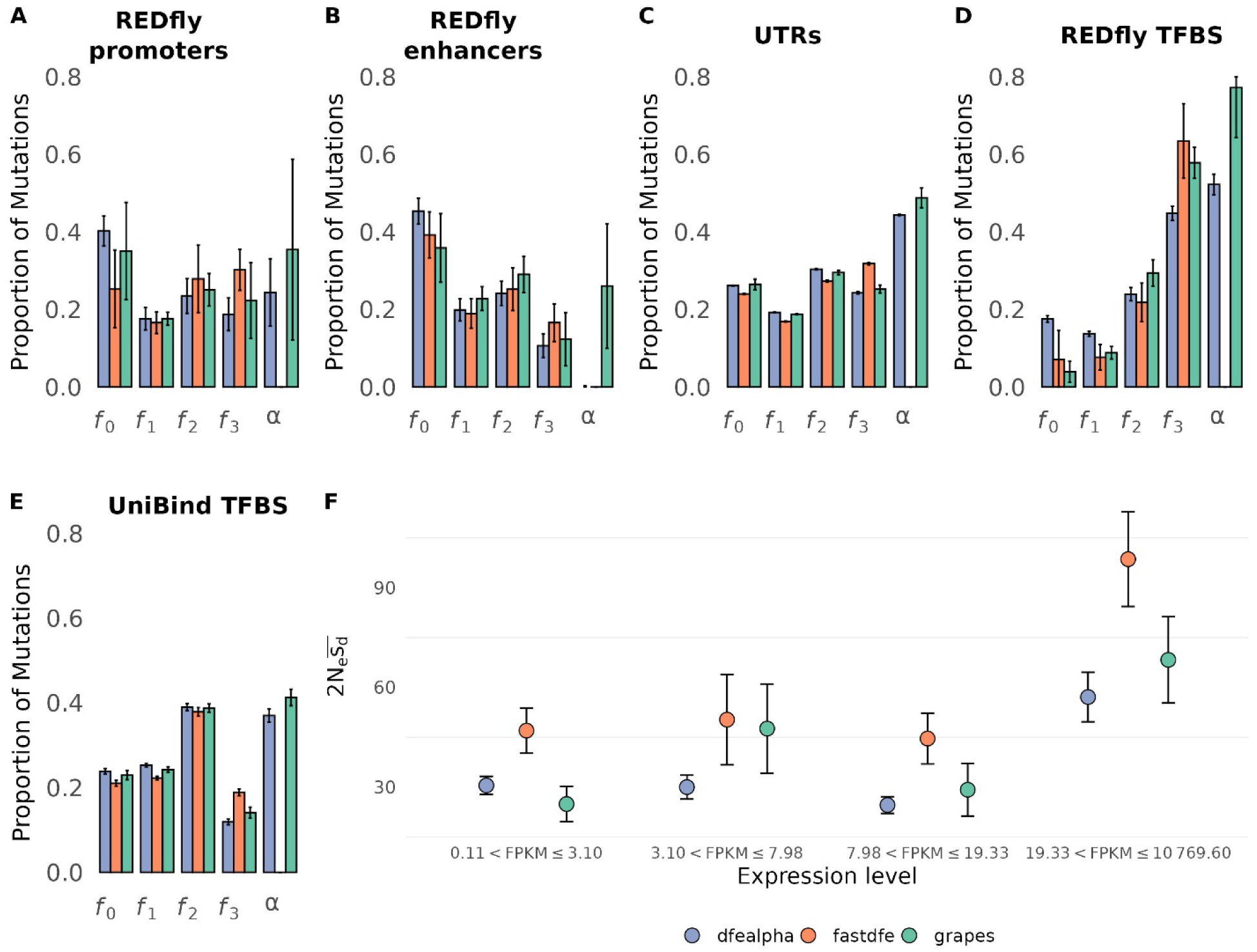
Inference of the distribution of fitness effects (DFE) of new mutations for (**A**) promoters, (**B**) enhancers, (**C**) UTRs, (**D**) transcription factor binding sites (TFBS) annotated by *REDfly*, and (**E**) TFBSs annotated by UniBind in the South population of *D. melanogaster*. The DFE is shown in terms of the proportion of mutations in effectively neutral (*f*_0_), weakly deleterious (*f*_1_), moderately deleterious (*f*_2_) and strongly deleterious (*f*_3_) classes of mutations, along with the proportion of substitutions that are beneficial (*α*). *α* predictions using *fastDFE* were excluded due to their inaccuracy in the benchmarking simulations. (**F**) The mean deleterious population scaled selection coefficient 2*N_e_s̅_d_* of the inferred DFE for UniBind TFBSs, stratified by the adult expression level of the nearest gene.

For promoters and enhancers, estimates of the proportion of beneficial substitutions varied considerably across subsampling replicates, with *GRAPES* consistently reporting higher values than *DFE-alpha* (Figure 5). This is likely due to a limited number of selected sites available for inference (∼18k for promoters and ∼50K for enhancers), which was lower than the threshold of 100,000 sites needed for accurate inference in our simulations. The mean estimate of the proportion of beneficial substitutions estimated by *DFE-alpha* was 0.25 in promoters, highly variable across subsampling replicates (-0.07 to 0.26) in enhancers, and 0.4 in UTRs (where ∼3.5 million sites were used for inference). For *REDfly* transcription factor binding sites, estimates of the proportion of beneficial substitutions were much higher than those for other non-coding regions for *DFE-alpha* (0.52) and *GRAPES* (0.77), though note again there were only ∼14,000 selected sites used for inference. Note that masking sites within repeats had no effect on our inference for specific non-coding regions (Figure S10).

### Selection on TFBSs in *D. melanogaster*

*REDfly* annotations are a small subset of all TFBSs in the genome, not intended to be representative of all TFBS. Thus, this dataset may be subject to an ascertainment bias, as they are hand-picked by researchers for experimental validation, often due to the interesting phenotypes they produce when knocked out, potentially biasing them towards the more strongly deleterious DFE and higher *α* estimates we observed. To understand if the DFE we observed in *REDfly* TFBS applies to a wider set of TFBS, we used the *UniBind* database (Puig et al. 2021), a computational annotation of TFBS based on ChIP-seq data. Instead of simply using ChIP-seq peaks to annotate TFBS, *UniBind* uses TFBS motifs from *JASPAR*, an open-source TF binding profile database (Fornes et al. 2020) to narrow down TFBSs to more realistic lengths. However, these annotations should not be interpreted as a complete set of functional TFBSs: some predicted sites may represent false positives, context-specific binding, or binding events to sites that do not measurably affect gene expression (see MacQuarrie et al. 2011). Overall, we obtained 258,618 selected sites and 1,069,601 neutral flanking sites from these regions as opposed to 14,022 and 25,111 used previously. In the South population, the inferred DFE for UniBind TFBS using *DFE-alpha* (2*N_e_s̅_d_* = –39; Figure 5E) was much less deleterious than the *REDfly* TFBSs 2*N_e_s̅_d_* = –387. Additionally, the proportion of beneficial substitutions was ∼0.4, much lower than the values obtained for the *REDfly* TFBSs. The higher proportion of beneficial substitutions inferred for REDfly TFBSs may reflect the small number of selected sites or ascertainment bias toward experimentally validated functional elements; however, the less deleterious DFE inferred for UniBind TFBSs may reflect false positives.

It is plausible TFBSs near highly expressed genes experience stronger purifying selection because these genes often require sustained high transcript/protein output (and are frequently constitutively expressed), such that cis-regulatory mutations that change gene expression are more likely to impose deleterious dosage deficits or imbalances (Gout et al. 2010; Makanae et al. 2013; Rice and McLysaght 2017). Consistent with this view, total gene expression has frequently been negatively associated with rates of protein sequence evolution (Pál et al. 2001; Subramanian and Kumar 2004; Larracuente et al. 2008; Gout et al. 2010) and with evolutionary rates in nearby regulatory regions (Dukler et al. 2022; S. Chen et al. 2024). Utilizing the *UniBind* dataset we find that the 2*N_e_s̅_d_* of TFBSs near genes with moderate to high expression (fragments per kilobase of transcript per million mapped reads, FPKM>19.33) is almost twice those of the TFBSs near genes with low to moderate expression (FPKM below 19.33; Figure 5F). This indicates that, while we consistently inferred a moderately deleterious DFE for different types of non-coding regions throughout the genome on average, the DFEs of specific non-coding regions may be dependent on their functional roles in the genome.

### Incorporating non-coding DFEs improves predictions of background selection across the *D. melanogaster* genome

Analytical methods to calculate *B*, the expected neutral diversity in the presence of purifying selection relative to expected diversity under neutrality, have been developed to predict genomic variation (Charlesworth et al. 1993; Hudson and Kaplan 1995; Nordborg et al. 1996; Comeron 2014; Elyashiv et al. 2016; Johri et al. 2020). Recently, these calculations have been made practical at genomic scale by *Bvalcalc* (Marsh et al. 2026), which provides an efficient implementation for computing regional and genome-wide *B*-maps while allowing annotation-specific DFEs, recombination rate maps, gene conversion, and a single change in population size. Applying these methods to all *D. melanogaster* autosomes, we find that incorporating non-coding DFEs substantially improves the prediction of diversity across the genome (Figure 6). Coding mutations alone yield a mean *B* of 0.84 (median 0.94) but including conserved non-coding and UTR mutations reduced these values to a mean of 0.64 (median 0.82), reflecting the large contributions of non-coding DNA to background selection. This adjustment markedly improves the fit of *B* maps to observed polymorphism (Pearson *R*^2^ at a 1Mb scale from 0.49 with coding regions only to 0.76 with UTRs and non-coding regions added; Figure S11).

**Figure 6:**
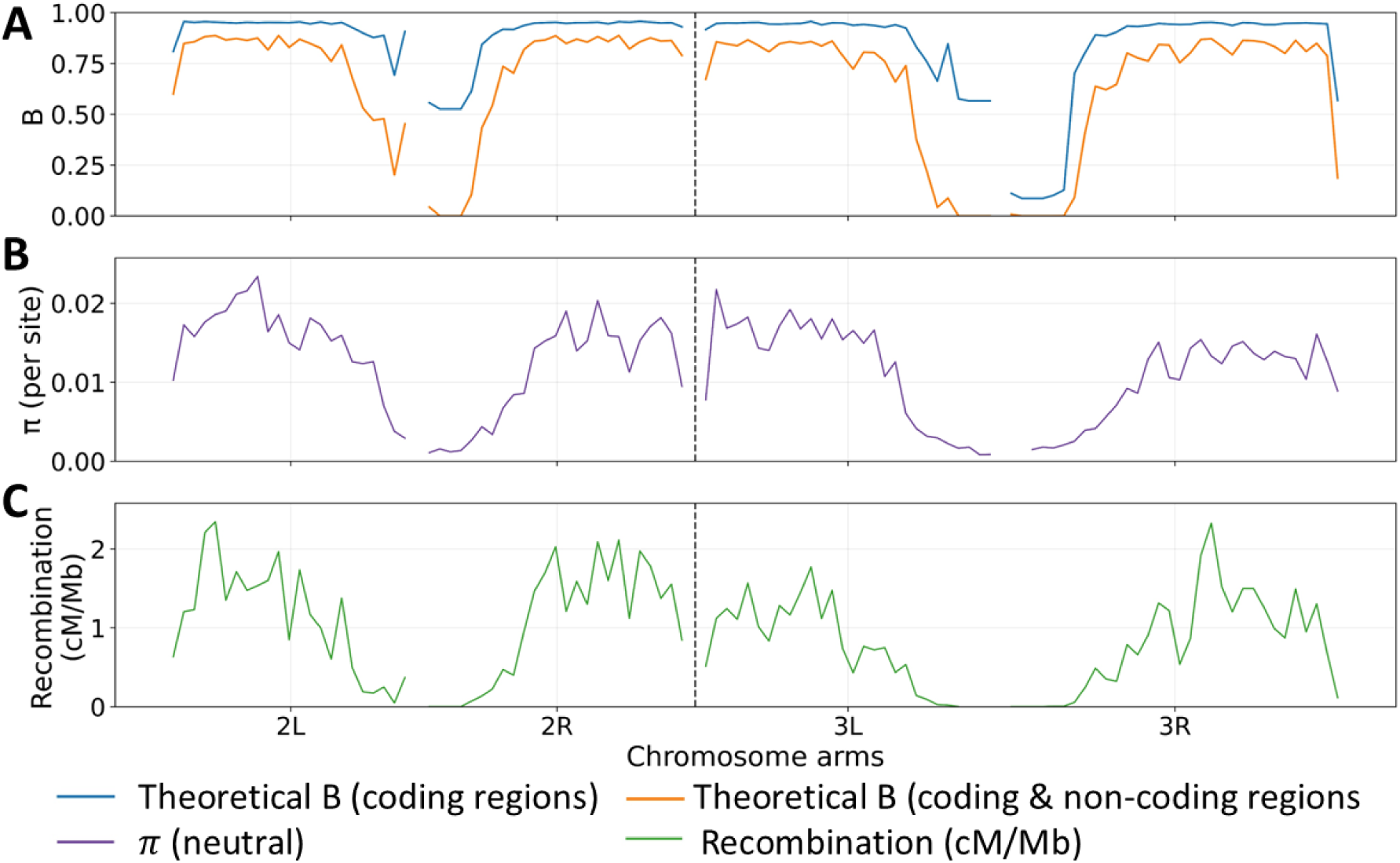
Comparison of predicted *B* maps in *D. melanogaster* with and without non-coding regulatory regions. (**A**) Predicted genome-wide *B* maps across autosomes of D. melanogaster calculated with *Bvalcalc*. Yellow lines depict *B* values calculated when only coding sequences (75% selected) are considered. Blue lines show the predicted values when UTRs and conserved non-coding *phastCons* elements were also considered. (**B**) Observed nucleotide diversity (*π*) at synonymous and putatively neutral intergenic sites in the South population of *D. melanogaster*. (**C**) Sex-averaged recombination rates across chromosome arms of *D. melanogaster* from Comeron (2002).

## DISCUSSION

### Comparison to previous estimates of DFE in non-coding regions in Drosophila

This study extends our understanding of the distribution of fitness effects (DFE) in *D. melanogaster* non-coding DNA by analyzing multiple classes of putatively functional annotations with a combination of DFE inference methods. The genomes used in this study include multiple *D. melanogaster* individuals from three distinct sub-Saharan African populations, including a rural population from Zimbabwe thought to be close to the ancestral range of the species (Coughlan et al. 2022). Our estimate of the deleterious DFE in conserved non-coding regions closely matches earlier results obtained using *DFE-alpha* on conserved non-coding regions from a small segment of the X-chromosome in 6 *D. melanogaster* individuals from Zimbabwe (Casillas et al. 2007): we infer a 2*N_e_s̅_a_* = –37.9 and a shape parameter *β* = 0.37 in our South population, compared with their 2*N_e_s̅_a_* = –61.4 and *β* = 0.31. Likewise, our estimates of the proportion of beneficial fixations within these conserved non-coding regions (0.27-0.31 in the South population) align closely with Andolfatto’s (2005) inference from conserved non-coding DNA (0.27 and 0.20, respectively, though Andolfatto’s data was all X-linked), although both studies diverge from the lower *α* (0.13) reported by Casillas *et al*. (2007), which was made using a combination of conserved and non-conserved non-coding DNA. As in Andolfatto (2005), our estimate of the proportion of beneficial fixations in UTRs (∼0.46 in South from *DFE-alpha*) was higher than that of other non-coding regions, though not as high as the value of 0.58 reported in that study.

Across our three sub-Saharan African populations, we found little evidence for systematic population-specific shifts in the deleterious DFE. While individual methods occasionally reported differences in 2*N_e_s̅_d_*, these were rarely reproduced across methods and annotations, consistent with broadly shared selective constraints on functional DNA. This may partly reflect the recency of population divergence and range expansion in *D. melanogaster*. Demographic modeling of African populations suggests the ancestors of the East and West populations split from a rural South population ∼ 500k generations ago, implying extensive shared standing variation and limited time for population-specific SFS distortions to accumulate due to the large *N_e_* of *D. melanogaster* populations (Sprengelmeyer et al. 2020).

### Evolution of non-coding vs coding regions

We found that the mean strength of purifying selection against deleterious mutations in 0-fold coding sites (2*N_e_s̅_d_*) was substantially larger than that in conserved and experimentally validated non-coding regions, differing by roughly an order of magnitude. Interestingly, similar contrasts between coding and conserved non-coding regions have been reported in *M. m. castaneus*, *H. sapiens*, and *C. grandiflora* (summarized in Table 2), despite variation in methods used to annotate conserved and putatively neutral sites and to calculate divergence. Estimates of the proportion of beneficial fixations are more difficult to obtain, but where available, this proportion is usually higher in coding regions than in conserved non-coding regions, with the exception of *C. grandiflora*, where coding *α* was 0.41 compared to 0.54 in conserved non-coding regions (Table 2). Expanding such analyses to a wider range of taxa will help determine whether these patterns represent a general feature of genome evolution.

To compare the contributions of new mutations at coding and non-coding regions to fitness, we use the DFEs estimated by *DFE-alpha* for the South population. Assuming equal per-site mutation rates, nonsynonymous coding sites (9,343,752 autosomal sites) accounted for only 12.0% of new deleterious mutations per diploid individual/ generation, compared with 78.2% from conserved non-coding regions (60,672,393 sites, comprised of phastCons elements not overlapping exons) and 9.7% from UTRs (7,529,622 sites). To estimate their relative contributions to deleterious fitness effects, we assumed multiplicative fitness effects across sites and calculated the expected reduction in fitness in one generation from newly arising mutations in a wild type background by integrating over the inferred gamma DFEs (see Methods). Under these assumptions, nonsynonymous mutations contributed 71.6% to reducing fitness per generation, compared with 21.7% from conserved non-coding regions and 6.6% from UTRs. Thus, although conserved non-coding regions generate more deleterious mutations each generation, coding mutations contribute more to aggregate fitness effects because they are subject to substantially stronger selection. This is qualitatively consistent with Booker *et al*. (2021), who found that protein-coding regions in mice could contribute more to adaptive fitness gains than conserved non-coding elements despite the greater frequency of non-coding beneficial substitutions. However, their analysis considered fitness gains from beneficial substitutions, incorporating fixation probabilities, whereas our calculation considers the immediate effects of new deleterious mutations.

. Similarly, estimates of the proportion of beneficial fixations were lower in non-coding regions relative to non-synonymous mutations but make up a larger proportion of beneficial fixations (69.7%) than non-synonymous mutations (16.0%) and UTRs (14.3%) when considering the large proportion of the genome they comprise. It is likely that the mean strength of fixations of beneficial mutations is much larger in coding vs non-coding regions, and because population-genetic methods are more likely to detect sweeps of strongly favorable mutations (Soni et al. 2023), the number of sweeps detected in coding regions might be similar or ever larger than those detected in non-coding regions. Inference of the DFE of beneficial mutations in non-coding regulatory regions would be needed to make such predictions. For instance Campos et al. (2017) estimated large population-scaled beneficial effects in *Drosophila* (nonsynonymous 2*N_e_s̅_a_* = 508 vs. UTR 2*N_e_s̅_a_* = 260), while Booker et al. (2021) inferred two classes of advantageous mutations in *M. m. castaneus* and found substantially stronger selective effects in exons than CNEs (2*N_e_s̅_a_* = 3100 and 105 for exons vs. 950 and 3.5 for conserved non-coding regions). Overall, our results suggest that non-coding regions contribute a majority of new deleterious mutations and beneficial substitutions in *D. melanogaster* populations.

### Evolution of TFBSs

Although we inferred highly similar DFEs for most non-coding annotation types in our high-confidence *REDfly* annotations, transcription factor binding sites (TFBS) were inferred to have a highly deleterious DFE with a large proportion of beneficial fixations. One plausible explanation is *cis*–*trans* compensatory coevolution, in which mildly deleterious substitutions in transcription factors alter binding or regulatory output and are subsequently compensated by changes in cognate TFBSs that restore function, making some TFBS substitutions appear beneficial in their derived genetic background rather than reflecting *de novo* adaptive optimization of the site (Landry et al. 2005; Kuo et al. 2010; Lynch and Hagner 2015). However, when we inferred the DFE for a larger set of TFBS from the *UniBind* database (Puig et al. 2021), the DFE was more moderately deleterious like other non-coding regions, raising doubts about whether our inference in the *REDfly* dataset applies to TFBS in general or is due to an ascertainment bias or using a small number of selected sites. On the other hand, *UniBind* identifies TFBSs computationally from genome-wide binding data without requiring experimental validation of regulatory function at each site. Consequently, some predicted sites may be false positives, while others may bind transcription factors without substantially affecting gene expression or fitness. Including such sites could weaken the inferred signal of purifying selection relative to the experimentally validated *REDfly* TFBSs. Thus, the contrast between *REDfly* and *UniBind* likely reflects both differences in the functional importance of TFBSs and their methods of identification..Future progress on high-quality annotations of TFBS will yield more insights into their evolution.

### Implications of the prevalence of mildly and moderately deleterious mutations in non-coding regions

Purifying selection at one site affects variation at linked alleles, and thus correctly modeling purifying selection is necessary to understand patterns of genetic variation across the genome, which is important for downstream tasks like demographic inference (Nordborg et al. 1996; Johri et al. 2021; Johri et al. 2022). Though the *B* maps we calculated should only be viewed as an approximate proof of concept (Figure 6), accurately modeling the contribution of non-coding deleterious mutations is clearly essential for interpreting nucleotide diversity in *D. melanogaster*, especially because a large number of new moderately deleterious mutations occur every generation at these sites, which persist long enough to exert substantial background selection effects on linked neutral sites as they are purged from the population (Nordborg et al. 1996). The excess of nearly neutral and mildly deleterious mutations in the noncoding DFE is also relevant in regions with very little recombination, as these alleles will be more likely to experience Hill-Robertson interference effects or associative overdominance relative to mutations under strong selection (Hill and Robertson 1966; McVean and Charlesworth 2000; Comeron and Kreitman 2002; Becher et al. 2020). Consequently, many non-coding deleterious alleles are expected to have elevated fixation probabilities in regions of little/no recombination, increasing mutation load and reducing fitness (Charlesworth and Jensen 2021).

### Caveats

Although the overall consistency of our findings is encouraging, several limitations must be considered. For our “high-confidence” set of functional non-coding regions, we utilized the *REDfly* database, which may be biased towards the annotation of non-coding regions under stronger selection due to the manual curation process, in which researchers may be more likely to pick regulatory regions that cause visible, large-effect phenotypic changes when altered. As confirmation of this, the more comprehensive computational annotations from the *UniBind* TFBS dataset were inferred to have a less deleterious DFE than the *REDfly* TFBS, though note the *UniBind* dataset itself may also be subject to ascertainment bias due to more representation of transcription factors (TFs) with a higher number of TFBS or stronger ChIP-Seq peaks (Puig et al. 2021; Tahara and Ozaki 2025). While it is encouraging that the high-confidence *REDfly* and conserved non-coding (*phastCons*) regions yield very similar DFEs, it is worth noting that *phastCons* elements are explicitly defined by reduced divergence, which biases against inclusion of very weakly selected sites. Despite a thorough filtering of our putatively neutral sites, some weakly selected sites may remain, as evidenced by smaller divergence values in neutral non-coding sites relative to synonymous sites and short introns (Table 1). However, synonymous sites in *Drosophila* can themselves experience selection, particularly associated with codon usage bias, although recent analyses suggest that most are effectively neutral (Akashi 1995; Jackson et al. 2017; Johri et al. 2020). Deflated divergence values for the neutral baseline will bias estimates of the proportion of beneficial substitutions upwards (Akashi 1995). Additionally, recent studies have shown that weak direct selection on the set of putatively neutral sites may cause bias in estimates of the DFE towards strongly deleterious mutations, though this bias was not pronounced when fewer than 20% of synonymous sites were under weak purifying selection (Zurita et al. 2025). Thus, if a small minority of noncoding sites are under direct purifying selection, our estimates of the DFE should be reasonably accurate.

Recent simulation work suggests that unmodeled fine-scale heterogeneity in mutation and recombination rates can inflate uncertainty and bias DFE inference, often toward an excess of effectively neutral mutations (Soni et al. 2024). This sensitivity is expected because most DFE methods implicitly assume that the neutral reference captures the same biases in the SFS caused by the local mutation rate and local effects of selection on linked sites as the selected sites. In that work, heterogeneity is imposed at a very fine scale by assigning each 1 kb segment a mutation and/or recombination rate drawn from a uniform distribution spanning broad bounds (mutation rate between 1 × 10^−9^ − 6.1 × 10^−9^ /site/generation and a recombination rate between 1.27 × 10^−9^ − 7.4 × 10^−8^ /site/generation). In contrast, our forward simulations assume empirically derived recombination-rate variation and mutation rate heterogeneity across the *D. melanogaster* genome, using neutral sites flanking selected non-coding regions as a neutral reference. Under these conditions, we find that DFE inference is generally accurate for DFEs primarily comprising moderately and strongly deleterious mutations, with a small amount of bias for DFEs dominated by mildly deleterious mutations, suggesting that the extent of misinference depends heavily on the genomic scale at which mutation rates significantly differ in organisms. Bias in DFEs comprising predominantly weakly deleterious mutations arose when beneficial mutations were present and increased as beneficial effects became stronger and rarer, with the product *f*_pos_ × 2*N_e_s*_pos_ held constant. This pattern suggests that reduction in effective population size due to sweeps may result in the inference of lower values of 2*N_e_s*_d_.

Alternatively, selective sweeps resulting from the fixation of strongly beneficial mutations may result in higher frequencies of deleterious mutations (compared to that expected under neutrality) due to genetic hitchhiking, thus affecting inference. A related concern is Hill–Robertson interference in regions of very low recombination, which can distort the SFS and make neutral and selected spectra more similar, biasing DFE inference (Daigle and Johri 2025). However, Daigle and Johri found that such interference produced appreciable mis-inference under the most extreme scenarios of little/no recombination, with error increasing sharply when *B* due to linked effects was strongly reduced below ∼0.5. Because the majority of the *D. melanogaster* genome is well above this threshold according to the *B* map inferred in this work (Figure 6), and consistent with the minimal bias observed in our *Drosophila*-like simulations, pervasive interference of this magnitude is unlikely to dominate genome-wide inference.

Estimating the proportion of beneficial substitutions is inherently challenging. Fluctuations in ancestral population size that may no longer be statistically detectable, determine the probability of fixation of mildly deleterious mutations and thus the proportion of beneficial substitutions (Eyre-Walker 2002). Similarly, DFE inference approaches that employ pairwise divergence assume that the outgroup population size is the same as the size of the focal species and that the estimate of *α* includes substitutions occurring on branches leading to both the outgroup and focal species. Thus population-size misspecification of ancestral and outgroup species can alter the expected fixation rate of weakly deleterious mutations and thus bias *α*. Zhen et al. (2021) re-estimated the proportion of beneficial fixations for the *D. melanogaster–D. simulans* pair under a complex demographic model with a large ancestral population size (*Nanc*∼1.143 × 10^7^), a *simulans* lineage maintaining the ancestral size after the split, and *D. melanogaster* growing from 2.79 × 10^6^ to 7.63 × 10^6^ (based on their earlier demographic estimates). Under these assumptions, their model produced a lower *α* (0.60) estimate than *DFE-alpha* (0.71) on the same dataOur simulations suggest that if the effective population size of *D. melanogaster* and *D. simulans* is 1 million and 1.5 million respectively, we would be overestimating *α* by ∼15.4%. Thus, our empirical estimates of *α* in coding and non-coding regions may be upwardly biased, although the magnitude of this bias remains uncertain and dependent on the true demographic histories of these species, and could be stronger if the difference in effective population sizes is larger than we assume here. Because we used only *D. simulans* as an outgroup, substitutions could not be assigned to either lineage. Additional outgroups would allow branch-specific divergence to be estimated and potentially reduce this source of bias. This suggests that the true values of *α* in coding and non-coding regions are likely 0.46 and 0.23, respectively. Further work is needed to compare the predictions of more complex demographic models in the ancestral population to standard approaches to determine the extent of expected bias.

Finally, as recent migration has been reported between African *D. melanogaster* populations (Kapopoulou et al. 2018; Coughlan et al. 2022), our pooling of samples in the East and West populations across geographic locations could alter the SFS in ways not captured by the simple demographic models used for DFE inference. However, no substructure within these populations was identified. Country-level analyses of samples from the East and West populations yielded similar DFEs at *phastCons* sites to those inferred when samples were grouped by geographic region (Figure S8). Moreover, recent simulation work suggests that DFE inference can be reasonably robust to moderate levels of unmodeled population structure (Gossmann et al. 2010; Daigle and Johri 2025). Other complexities such as the effects polymorphic inversions (Corbett-Detig and Hartl 2012; Kapun and Flatt 2019) on inference of the DFE merit further investigation.

## METHODS

### Identification of putatively selected and neutral regions

Experimentally validated regulatory region annotations were derived from from the *REDfly* project (Keränen et al. 2022; *v9.6.4*), which curates regulatory elements supported by experimental approaches including reporter assays, ChIP-seq, *DNase I* hypersensitivity assays, electrophoretic mobility shift assays, and related functional assays. Annotation data for microRNA (miRNA) and miRNA primary transcripts was downloaded directly from *miRBase* (accessed Feb 2024, https://www.mirbase.org/download/; Kozomara et al. 2019). These were termed “high-confidence” regions due to their experimental validation. *phastCons* scores for evolutionary conservation of sites and genomic elements (Siepel et al. 2005) were also downloaded from the UCSC Genome Browser (accessed Jan 2024, https://genome.ucsc.edu/cgi-bin/hgGateway). To obtain other potentially functional regions in the non-coding regions of the genome, the *D. melanogaster* r6.56 annotations were downloaded from *FlyBase* (accessed Mar 2024, https://ftp.flybase.net/releases/FB2024_01/dmel_r6.56/gff/; Öztürk-Çolak et al. 2024) and the coordinates and annotations of regulatory regions from the *ORegAnno* 3.0 database (Lesurf et al. 2016) were downloaded using the table browser tool in the UCSC Genome Browser (accessed Jan 2024, https://genome.ucsc.edu/cgi-bin/hgGateway; Raney et al. 2024). *FlyBase* annotations are derived from curated gene models supported by transcriptomic data, sequence homology, computational predictions, and published experimental evidence. *ORegAnno* compiles regulatory annotations from published studies and external databases, incorporating experimental evidence of transcription factor binding and regulatory activity (Lesurf et al. 2016; Öztürk-Çolak et al. 2024). The annotations from *FlyBase* and *OregAnno* were filtered to retain only relevant labels (Table S3), which were focused on potentially functional annotations provided by *FlyBase*, which we termed “low-confidence regions”. Coding regions (CDS), untranslated regions (UTRs), high-confidence regions, *phastCons* elements, and low-confidence regions comprised the final non-overlapping and exhaustive set of regions considered to comprise the genome. While CDS, UTRs, high-confidence regions, and *phastCons* elements were considered to be under direct selection, the purpose of annotating low-confidence regions was primarily to exclude non-coding sites that may be considered putatively neutral. BED files summarizing downloaded locations and data sources of CDS, UTR, *phastCons*, low- and high-confidence, and neutral regions can be found at https://doi.org/10.5281/zenodo.18777928 in the folder nonoverlapping_annotations.zip.

A non-overlapping annotation of the *D. melanogaster* genome was constructed as follows. First, any region overlapping a CDS annotated in the *FlyBase* r6.56 GFF was defined as coding. UTRs were annotated next, with any UTR region also annotated as CDS being annotated as coding instead. High-confidence regulatory regions were defined next, excluding overlaps with CDS and UTRs. Additionally, sub-annotations were created depending on the annotation label of the high-confidence regions (i.e., promoter, enhancer, and transcription factor binding site (TFBS)). Next, *phastCons* elements were annotated (where regions overlapping CDS, UTR or high-confidence regions were excluded), followed by remaining low-confidence computational annotations listed in Table S3, again excluding previously annotated regions (i.e. no low-confidence element overlapped a high-confidence one or a UTR).

All remaining sites not overlapping any of these annotations were considered putatively neutral non-coding sites. We further filtered these neutral sites using the downloaded *phastCons* per-site scores, which represent the posterior probability that a site belongs to an evolutionarily conserved state (Siepel et al. 2005). Per-site *phastCons* scores were strongly bimodal, with the retained neutral sites concentrated near zero relative to functional coding and non-coding regions (Figure S12A). More stringent thresholds progressively increased nucleotide diversity and divergence, as expected from removing conserved sites, but substantially reduced the number of neutral sites available for inference. We therefore selected 0.1 as a balance between these factors (Figure S12C). A more stringent cutoff of 0.05 produced only modest changes in divergence and DFE estimates and did not enrich for high-divergence outliers (Figure S12B and D). Non-coding neutral sites within 5 kb of high-confidence regions and *phastCons* elements were annotated as flanking neutral sites for DFE inference. Additional sets of flanking neutral annotations were generated specifically for selected subregions (promoters, enhancers, and transcription factor binding sites) within the high-confidence annotations and for UTRs. The final set of putatively non-coding neutral sites identified have been provided at https://doi.org/10.5281/zenodo.18777928 in the folder nonoverlapping_annotations as the file neutral_intergenic.bed.

We obtained a set of computationally predicted TFBSs for *D. melanogaster* from the *UniBind* database (Puig et al. 2021; accessed May 2025; https://unibind.uio.no/downloads/). This resource provides robustly filtered TFBS predictions derived from multiple ChIP-seq datasets, reducing false positives and standardizing TFBS size across experiments. UniBind TFBSs were analyzed separately and were not incorporated into our high-confidence genome annotation because, unlike the individually curated experimentally validated REDfly elements, these sites are computationally inferred from genome-wide TF-binding datasets. Instead, we filtered the *UniBind* TFBSs to exclude overlaps with CDS and UTRs, while allowing overlaps with other non-coding annotation categories, including high-confidence, *phastCons* and low-confidence regions, and drew their flanking neutral sites from the pre-defined filtered neutral set. Repeat annotations were obtained from the UCSC Genome Browser Table Browser (RepeatMasker track, accessed October 2025) for the *D. melanogaster* dm6 assembly (Karolchik et al. 2004). When all monomorphic and polymorphic sites within repeats were filtered (∼10% of sites), we obtained very similar parameter values of the DFE estimated for each category of non-coding sites (Figure S10).

### Obtaining the site frequency spectra (SFS) from population genomic data

We obtained single nucleotide polymorphism (SNP) data for chromosomes 2 and 3 of *Drosophila melanogaster* from the VCF file provided by Coughlan *et al*. (2022). Details on sequencing and variant calling procedures are described in the supplementary methods of their publication. Briefly, their provided VCF was filtered to remove indels, retaining only invariant and biallelic SNP sites. Quality filters included a minimum variant quality score of 30, minimum sequencing coverage of 5×, minimum genotype quality score of 30, and a maximum per-site missingness threshold of 25% (*i.e.*, sites missing genotype calls in >25% of individuals were excluded). Additionally, genomic regions within 100kb of nine major polymorphic inversions were removed from this VCF for all individuals in the population. To select samples for downstream analyses, we used the ancestry assignments reported by Coughlan *et al*. (2022). We retained only individuals whose geographic sampling locations were concordant with their assigned ancestry clusters and grouped them into three focal populations (“South1”, “East,” and “West”) following Coughlan et al.’s labels (see Figure 1 in Coughlan *et al*. (2022)). Our final “South” population included only individuals newly sampled from Mana Pools National Park in Zimbabwe; the “East” population comprised samples from Rwanda and Uganda; and the “West” population contained samples from Cameroon, Gabon, Ghana, Guinea, and Nigeria. Sample GAB187, which displayed unusually elevated heterozygosity, was removed. We also retained 13 *D. simulans* individuals to identify outgroup polymorphism and excluded sites that were fixed in the focal *D. melanogaster* population but polymorphic in *D. simulans*, preventing such sites from being ambiguously classified as fixed differences between the species. We provide the final VCF file at https://doi.org/10.5281/zenodo.18777928 in the folder vcf_for_dfereg_simulansfilter.noRepeats.

The sequencing design differed among populations. Genomes from the South populations were derived from diploid isolines and were analyzed as diploid samples (141 total). In contrast, the East and rural West genomes were generated from haploid embryos but represented as diploid genotypes in the VCF. We therefore recoded these samples as haploid before constructing the SFS: homozygous-reference and homozygous-alternate genotypes were converted to single reference or alternate alleles, respectively, whereas heterozygous and missing genotypes were treated as missing. Heterozygous calls were not randomly resolved because in true haploid embryos such calls are more likely to reflect mapping, copy-number, or genotype-calling error. The resulting datasets contained 29 East and 26 West haploid genomes.

We used the parsing tools provided by *fastDFE* (Sendrowski and Bataillon 2024) to calculate the SFS for each population. We inferred the ancestral state using the algorithm within *fastDFE* for each site in our VCF, which included polymorphisms and invariant sites. We used the *AdaptivePolarizationPrior*, which is similar to the widely used program *EST_SFS* (Keightley and Jackson 2018) allowing divergence with the setting *allow_divergence=True*. Briefly, the polarization method used a probabilistic nucleotide-substitution framework to estimate the most likely ancestral allele at each site, accounting for uncertainty in cases where outgroup data are missing or ambiguous, which substantially reduces misinference compared to simple parsimony approaches (Keightley and Jackson 2018). Because this model estimates polarization probabilities from a fixed number of ingroup samples, we set *n*_*ingroups* = 21 as recommended in the *fastDFE* documentation, meaning that sites were used for ancestral-state inference only when at least 21 ingroup samples were present and that the ingroup data were randomly subsampled to this constant size for estimating the polarization prior. The maximum number of sites (max_sites) to estimate per-branch substitution rates and to calibrate the polarization prior was set at 200,000 as recommended in the *fastDFE* documentation. A single *D. simulans* individual, DRM18, was used as the outgroup for ancestral-state inference. The folded and unfolded SFS for our annotated genomic regions were calculated using the *fastDFE* VCF parser, using the previously polarized VCF as input (sites that were not successfully polarized were excluded from further analysis). We used *fastDFE*’s built-in codon-degeneracy parser (default settings) to classify coding sites as 0-, 2-, 3-, or 4-fold degenerate; only 0-fold and 4-fold sites were used for DFE inference. Site annotation was based on the *D. melanogaster FlyBase* r6.56 GFF together with the corresponding *RefSeq* genome assembly (GCF_000001215.4; accessed May 2024). These annotations were then used to construct the SFS for 4-fold and 0-fold degenerate sites.

### Calculation of summary statistics using putatively neutral sites

Population-genetic summary statistics were calculated from the unfolded site-frequency spectrum (SFS) using a custom *R* script. Nucleotide diversity (*π*) was computed as 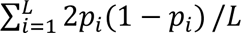 where *L* is the total number of sites and *p_i_* is the frequency of the derived allele at site *i*. Watterson’s theta (*θ*_W_) was computed from the number of segregating sites (*S*) using 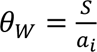 (with 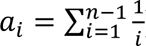), where *n* is the sample size, and was reported per site by dividing by the total number of sites (Watterson 1975). Tajima’s *D* was calculated from the difference between *π* and *θ*_W_, standardized by its expected variance using the sample-size-dependent constants *e*₁ and *e*_2_ as defined by Tajima ( 1989). Finally, divergence was computed as the fraction of monomorphic sites that are fixed differences (using the bin of the unfolded SFS that corresponds to a frequency of 1). Pairwise *F*_ST_ was calculated for 4-fold degenerate sites in our filtered VCF using the Weir and Cockerham(1984) estimator implemented in *scikit-allel* (version 1.3.13; Miles et al. 2024). To estimate uncertainty, we used a block-jackknife approach, dividing the genome into non-overlapping 1-Mb blocks and recalculating genome-wide *F*_ST_ after excluding each block in turn. Jackknife standard errors were calculated from these leave-one-block-out estimates and used to construct 95% confidence intervals and one-sided *p*-values testing whether *F*_ST_ was greater than zero. This procedure was applied to comparisons among the three focal populations, among geographic sampling locations within populations, and between inversion-genotype classes. For analyses comparing individuals with different inversion genotypes, individuals with missing or heterozygous inversion genotypes in Coughlan et al. (2022) were excluded.

### Forward-in-time simulations of *D. melanogaster* populations

Forward-in-time population genetic simulations were performed using SLiM 4.0.1 (Haller and Messer 2023). Population-genetic parameters were drawn from previous empirical estimates, including an ancestral population size of 1,000,000 prior to rescaling (Keightley et al. 2014; Sprengelmeyer et al. 2020). To make simulations computationally tractable, all simulations were rescaled linearly: employing best practices as suggested by Marsh (2026), population sizes and the number of generations were reduced, by a factor of 100, while mutation and recombination rates as well as the gene conversion initiation rate were multiplied by the same factor (Comeron and Kreitman 2002; Hoggart et al. 2007; Kim and Wiehe 2009; Uricchio and Hernandez 2014). Selection coefficients were likewise multiplied by the same factor to maintain the appropriate population-scaled selection strength. Simulations were run for 140,000 generations at this constant population size to allow for a 10*N_scaled_* generation burn-in and 4*N_scaled_* additional generations, where *N_scaled_* represents the population size after rescaling (10,000). Note that because population genetic approaches can only infer the distribution of population-scaled selection coefficients, it is important to maintain the same value of 2*N*_e_*s*_d_ after rescaling. The absolute value of *N*_e_ is not relevant unless the simulated population size, *N_scaled_*, is so small that it violates the assumptions of the diffusion theory (Johri et al. 2026).

As a realistic whole-genome simulation that could be completed in a reasonable time frame is likely to lead to biases due to rescaling (Marsh et al. 2026), including altered recombination dynamics, we simulated non-overlapping 100 kb windows of the autosomes, each of which was simulated independently with its own recombination rate (Comeron et al. 2012), mutation rate, and annotations matching that of the corresponding genomic region. We simulated five annotation classes: CDS, high-confidence regulatory, *phastCons* elements, low-confidence regulatory, and neutral. The deleterious DFE in CDS regions was modelled with four non-overlapping uniform distributions representing effectively neutral (0 ≤ 2*N_e_s_d_* < 1), weakly deleterious (1 ≤ 2*N_e_s_d_* < 10), moderately deleterious (10 ≤ 2*N_e_s_d_* < 100), and strongly deleterious (100 ≤ 2*N_e_s_d_*) mutations, comprising 25%, 49%, 4%, and 22% of all selected mutations, respectively, as inferred by Johri et al. (2020). The DFE in *phastCons* elements and low-confidence regulatory regions was modelled with 57% of new mutations kept strictly neutral while 43% of new mutations drawn from the same DFE modelled in coding regions; this was done so that 45% of genome-wide sites experienced selection, which is consistent with empirical estimates (Siepel et al. 2005), yielding 67.8%, 21.1%, 1.7%, and 9.5% of mutations in the effectively neutral, weakly deleterious, moderately deleterious, and strongly deleterious classes, respectively. In neutral regions, we assumed *s*=0 for all mutations. A gamma-distributed deleterious DFE with two parameters, mean (*s̅_d_*) and shape (*β*) were assumed. The exonic DFE was assumed to comprise four nonoverlapping uniform distributions because it directly reflected the empirical estimates of Johri et al. (2020), whereas the high-confidence regulatory DFE was modeled as a gamma distribution because these were the focal sites for which we tested DFE inference, and the gamma distribution is explicitly modeled by each of the inference methods evaluated here. We simulated DFEs with predominantly weakly deleterious (2*N_e_s̅_d_* = 5 and *β* = 0.9), moderately deleterious (2*N_e_s̅_d_* = 50 and *β* = 0.5), or strongly deleterious mutations (2*N_e_s̅_d_* = 1000 and *β* = 0.3). When simulating positive selection, beneficial mutations were assumed to be 0.02% of new selected mutations in all non-neutral regions, with semi-dominant beneficial selective effects following an exponential DFE with 2*N_scaled_ s̅* = 250, reflecting parameters estimated by Campos *et al*. (2017). For each DFE, we completed simulations comprising the entirety of Chromosomes 2 and 3, which represented 963 100 kb regions for each scenario. 200 diploid individuals were sampled at the end of each simulation and used for inference.

The crossover rates for each 100 kb region were taken from the rates reported by Comeron *et al*. (2012) based on the *D. melanogaster* recombination rate calculator v2.30 (Fiston-Lavier et al. 2010) and halved to represent sex-averaged estimates. Gene conversion was modelled according to parameters estimated by Miller et al. (2016): the gene conversion rate prior to rescaling was set to *g_c_* = 1 × 10^−8^ (uniform across all bins) and the mean tract length was set to 440 bp, with only simple conversions modelled. The mean mutation rate across all regions was 3 × 10^−9^ per site/generation (Keightley et al. 2014) prior to rescaling and was varied in each 100 kb bin. Following Marsh and Johri (2024), we modeled regional mutation-rate heterogeneity using normal distributions parameterized by the empirical variation in synonymous divergence between *D. melanogaster* and *D. simulans* reported by Mackay et al. (2012). Because divergence at putatively neutral sites reflects the accumulation of mutations over evolutionary time, we used its coefficient of variation as a proxy for spatial variation in mutation rates. The mutation rates in each 100 kb region were drawn from normal distributions for each chromosome arm with a coefficient of variation of 0.403, 0.479, 0.576, and 1.022, corresponding to the coefficient of variation for divergence at 4-fold degenerate sites between *D. simulans* and *D. melanogaster* in the middle section of contigs 2L, 2R, 3L, and 3R, respectively (Mackay et al. 2012). Mutation rates were limited to between 0 and 6×10^-9^ per site/generation (twice the mean mutation rate), and values drawn outside of these ranges were set to the mean mutation rate. The recombination and mutation rate maps are available at https://doi.org/10.5281/zenodo.18777928 in the folder simulations/inputs.

### Simulating the outgroup and ancestral population

To evaluate the performance of DFE inference methods under alternative histories of the ancestral population, forward simulations were performed with a 200 kb genome, and a constant mutation and recombination rate of 3 × 10^−9^, 3 × 10^−8^ per base/generation respectively. Three mutation classes were included: neutral (*s* = 0), deleterious (gamma-distributed, 2*N_e_s̅_d_* = –1000, shape parameter 0.3), and beneficial (exponentially distributed, 2*N_e_s̅_a_* = 250, with relative genomic proportions of 50%, 49.98%, and 0.02%, respectively, such that 50% of sites experienced selection.

The ancestral population size was varied across runs (500,000; 1,000,000; or 2,000,000). After 20*N* generations of burn-in, this population split into an ingroup of 1,000,000 individuals and an outgroup of 1,500,000 individuals, mimicking the ∼50% larger effective population size of *D. simulans* relative to *D. melanogaster* (Andolfatto et al. 2011). An additional 25*N* generations were simulated to mimic the estimated divergence time of the two species. These values describe the unscaled demographic model. To reduce computational costs, simulations were run using a 200-fold rescaling factor in which population sizes and number of generations were divided by 200 and mutation rates, recombination rates, and selection coefficients were multiplied by 200. Fixations in either population occurring after the split were obtained, reflecting branch-specific substitutions. 10 replicates were simulated for each parameter combination and for each replicate, 200 genomes were sampled from the ingroup population. Unfolded selected and neutral SFS were constructed and used as input for *DFE-alpha*, *GRAPES*, and *fastDFE* which was used to estimate the deleterious DFE and the proportion of beneficial fixations.

### Obtaining the SFS from simulated data

Folded and unfolded SFS for high-confidence genomic regions and flanking neutral sites were computed using the parser provided by *fastDFE*. Across all analyses, we used the same filtered genomic regions as in the empirical inference to ensure the arrangement of selected and neutral sites was maintained. For flanking neutral sites specifically, we applied the same single-site *phastCons* filtering used in the empirical pipeline, thereby preserving the identical arrangement of neutral sites surrounding each high-confidence region.

### DFE inference using *fastDFE*, *GRAPES*, and *DFE-alpha*

We inferred the DFE using three methods: *DFE-alpha* (Keightley and Eyre-Walker 2007), *GRAPES* (Galtier 2016), and *fastDFE* (Sendrowski and Bataillon 2024). In all cases, deleterious fitness effects were assumed to follow a gamma distribution. For *DFE-alpha*, we first fit a two-epoch demographic model to the neutral SFS and then estimated the mean and shape of the deleterious DFE from the selected SFS conditional on the inferred demographic parameters; no beneficial component was included in this DFE fit.. *GRAPES* and *fastDFE* were run using folded and unfolded SFS as input assuming a gamma-distributed deleterious DFE and an exponentially distributed beneficial DFE. For analyses using the unfolded SFS, we enabled the built-in ancestral-state misidentification correction in *GRAPES* and *fastDFE*, which estimates a polarization-error parameter and applies it to both the neutral and selected SFS.

The proportion of beneficial substitutions (*α*) is not a parameter fitted directly by these methods, but is instead calculated from other inferred quantities, with the calculation differing among programs. For *DFE-alpha*, we used *est_alpha_omega* to estimate *α* post hoc from the excess selected divergence relative to that expected from the inferred deleterious DFE and neutral divergence (Eyre-Walker and Keightley 2009); divergence was therefore not used to fit the deleterious DFE itself. *GRAPES* was run using the GammaExpo model under its default [+A] formulation, which jointly fits polymorphism and divergence data. The proportion of beneficial substitutions was calculated as the excess of observed selected divergence over that expected from non beneficial mutations under the fitted DFE, using the default *N_e_s_a_*=5 threshold to distinguish beneficial from non beneficial substitutions. In the version of *fastDFE* used here, *α* was instead calculated from the substitution probabilities implied by the fitted gamma-plus-exponential DFE and did not use observed divergence. Thus, the *DFE-alpha* and *GRAPES* estimates incorporate divergence, whereas the *fastDFE* estimate represents the proportion of substitutions predicted to arise from the inferred beneficial DFE.

When performing inference in simulated regions, we generated replicate SFS by randomly subsampling sites without replacement from the full SFS (monomorphic and polymorphic sites), for varying numbers of individuals (25, 50, 100, and 200) and sites (1000, 5000, 10000, and 100000 selected sites). We kept the ratio of selected to neutral flanking sites constant when subsampling, equal to ratios used in empirical inference (Table S2). When performing inference using empirical data, we similarly subsampled approximately 75% of the available sites for each annotation without replacement to generate replicate SFSs for DFE inference, while keeping the ratio of selected to neutral sites constant.

### Assigning UniBind transcription factor binding sites to the nearest gene

Adult-male and female gene expression values were taken from *FlyAtlas2* (Leader et al. 2018; Krause et al. 2022). *FlyAtlas2* quantifies RNA-seq expression (reported as FPKM) separately for male and female adult tissues and larvae as well as whole-body expression values. Each UniBind TFBS was linked to the nearest *FlyBase* gene annotation. For equidistant ties, we selected the gene with higher adult-male mean FPKM. The linked gene inherited the adult male expression value above; genes were ranked by the adult male value and partitioned into four equal-sized quantiles (Q1–Q4). We used adult male expression to reduce the potential influence of ovary- and oogenesis-associated transcription on whole-body expression estimates (Church et al. 2023). Genes with adult male or female mean FPKM < 1 were excluded prior to analysis, to remove transcripts that are likely biologically negligible or with strong sex-specific expression levels.

### Calculating *B* maps

We computed expected levels of background selection across the *D. melanogaster* genome by obtaining values of *B*, the expected neutral nucleotide diversity under purifying selection relative to neutrality, using *Bvalcalc* (v1.3.0; https://github.com/JohriLab/Bvalcalc; Marsh et al. 2026). Recombination rates were taken from Comeron *et al*. (2012) and halved to represent sex-averaged estimates. The point mutation rate was set to 3 × 10^−9^per site/generation (Keightley et al. 2014). We ran *Bvalcalc* using a the single step-change demographic model parameterized using the effective population sizes inferred in Johri et al. (2020), which predict an ancestral *D. melanogaster* population size of 1,225,393 increasing to a current size of 1,347,932 at 1,600,000 generations ago. Finally, we generated two sets of *B* maps that differed only in the selected annotations and their associated DFEs: (i) coding annotations only, and (ii) coding, UTR, and *phastCons*-conserved non-coding annotations. Genome-wide *B* values, which were output in intervals of 1kb, were averaged to 1 Mb windows for downstream analyses. For each annotation class, we used the gamma-distributed DFEs inferred in this study by *DFE-alpha* for the South population: coding 2*N_e_s̅_d_* = 811, *β* = 0.347; UTR 2*N_e_s̅_d_* = 93.1, *β* = 0.24; and *phastCons* 2*N_e_s̅_d_* = 37.9, *β* = 0.371. Genome-wide *B* values were averaged in non-overlapping 1 Mb windows for downstream analyses. We estimated *π* from the South population in sliding windows with *scikit-allel*, using putatively neutral intergenic sites (excluding UTR, *phastCons*, low-confidence, and coding intervals) together with fourfold-degenerate synonymous sites within coding regions. To quantify fit, we calculated Pearson *R*^2^and Spearman *ρ* between mean *π* in 1Mb windows and the mean *B* averaged over the same windows, excluding any windows without neutral sites.

### Calculating expected loss of fitness from new mutations per generation

To compare the contributions of coding, conserved non-coding, and UTR sites to deleterious mutational input per generation, we assumed equal per-site mutation rates, semidominant mutations (ℎ = 0.5), independently drawn mutational effects, and multiplicative fitness across sites. For a diploid offspring carrying *M* new mutations with homozygous selection coefficients *s_i_* < 0, where *s_i_* is the selection coefficient of mutation *i*, relative fitness was defined as

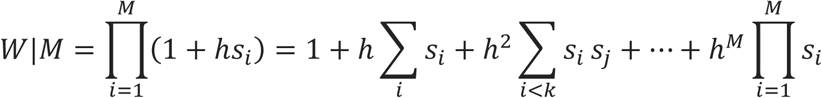

Then, the expected value of fitness would be given by:

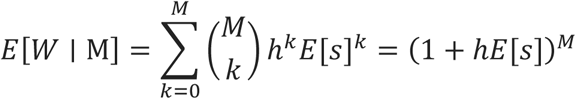

The first equality follows because the selection coefficients of distinct mutations belong to independent and identically distributed probability distributions. Thus, for any set of *k* distinct mutations, *E*[*s*_1_*s*_2_ ⋯ *s_k_*] = *E*[*s*_1_]*E*[*s*_2_] ⋯ *E*[*s_k_*]. The second equality follows from the binomial theorem.

Assuming that the number of new mutations per diploid offspring in category *i* is Poisson distributed, with *L_i_* mutable sites and per-site mutation rate *μ*, the expected number of mutations in a diploid genome (U) would be 2*L_i_μ*. Averaging over mutation counts, we have

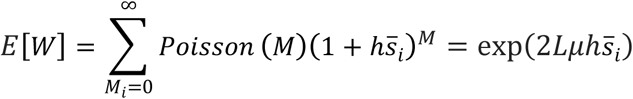

The expected reduction in offspring fitness relative to an otherwise identical offspring without new deleterious mutations was therefore

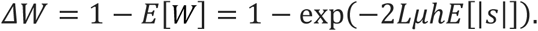

We converted the *DFE-alpha* estimate of the mean population-scaled deleterious effect, *x* = *E*[|2*N_e_s*|] and assumed *N_e_* = 10^6^ and *μ* = 3 × 10^−9^ per site per generation. Each category’s relative contribution was calculated as 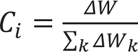.

## ACKNOWLEDGEMENTS

The research in this study was conducted using computational resources provided by ITS Research Computing at the University of North Carolina at Chapel Hill. Research reported in this publication was supported by the National Institute of General Medical Sciences of the National Institutes of Health under award number R35GM154969 to P.J. A.D. received additional support from NIH award R35GM13826 to Dan Schrider and NIGMS predoctoral training grant 5T32GM067553 during 2023-2024. The funders had no role in study design, data collection and analysis, decision to publish, or preparation of the manuscript. The authors declare no conflicts of interest.

## DATA AVAILABILITY

All data and code underlying this study are publicly available. The complete analysis workflow and scripts required to reproduce the empirical analyses and figures are available in the GitHub repository https://github.com/JohriLab/dfe_reg_regions. A permanent archive on Zenodo (DOI: https://doi.org/10.5281/zenodo.18777928) provides: (i) BED files specifying the genomic coordinates of exonic, *phastCons*, low- and high-confidence regulatory, and putatively neutral intergenic regions used for inference; (ii) the final filtered VCF for the three study populations used in the empirical analyses, together with the corresponding final accession list; and (iii) all code required to reproduce the simulations reported here (including the mutation and recombination rate maps), along with the simulation outputs analyzed in the manuscript. All parameter estimates from the empirical DFE analyses are reported in Tables S4 and S5.

## SUPPORTING INFORMATION

**Supplementary Figure 1:** The effects of the number of selected sites on the performance of *DFE-alpha*, *fastDFE*, and *GRAPES* in estimating the DFE of simulated high-confidence regulatory regions, when no beneficial mutations were simulated and the folded SFS was used as input. All inferences were performed with 200 individuals. Comparisons of the simulated and inferred DFE with a varying number of sites are shown in terms of the proportion of mutations in effectively neutral (*f*_0_), weakly deleterious (*f*_1_), moderately deleterious (*f*_2_), and strongly deleterious (*f*_3_) classes of mutations, along with the proportion of substitutions that are beneficial (*α*). The simulated DFE is denoted with grey bars, and the inferred DFE is shown as circles. In all panels, the error bars denote the standard deviation of proportions estimated from 10 replicates where sites were drawn with replacement from the total number of simulated sites.

**Supplementary Figure 2:** The effects of strength and frequency of beneficial mutations on the performance of *DFE-alpha*, *fastDFE*, and *GRAPES* in estimating the DFE of simulated high-confidence regulatory regions, when no beneficial mutations were simulated and the folded SFS was used as input. All inferences were performed with 200 individuals. Comparisons of the simulated and inferred DFE with a varying number of sites are shown in terms of the proportion of mutations in effectively neutral (*f*_0_), weakly deleterious (*f*_1_), moderately deleterious (*f*_2_), and strongly deleterious (*f*_3_) classes of mutations, along with the proportion of substitutions that are beneficial (*α*). The simulated DFE is denoted with grey bars, and the inferred DFE is shown as circles. In all panels, the error bars denote the standard deviation of proportions estimated from 10 replicates where sites were drawn with replacement from the total number of simulated sites.

**Supplementary Figure 3:** Effects of sample size on the performance of *DFE-alpha*, *fastDFE*, and *GRAPES* on estimating the (**A**) mean (*γ*) and (**B**) shape (*β*) of the *γ*-distributed deleterious DFE, along with the (**C)** proportion of selected substitutions that were beneficial (*α*) in simulated non-coding regions, using the folded SFS as input. Dotted lines represent the simulated parameters for *γ* and *β*, and the true value of *α* or the full set of simulated high-confidence regions.

**Supplementary Figure 4:** Effects of varying the number of individuals on the performance of *DFE-alpha*, *fastDFE*, and *GRAPES* when no beneficial mutations were simulated and the folded SFS was used as input. Shown here are estimates of the (**A**) mean (*γ*) and (**B**) shape (*β*) of the *γ*-distributed deleterious DFE, along with the (**C)** proportion of selected substitutions that were beneficial (*α*) in simulated non-coding regions. In all panels, the error bars denote the standard deviation estimated from 10 independent replicates.

**Supplementary Figure 5:** Using the unfolded SFS, effects of varying the number of individuals on the performance of *DFE-alpha*, *fastDFE*, and *GRAPES* on estimating the (**A**) mean (*γ*) and (**B**) shape (*β*) of the *γ*-distributed deleterious DFE, along with the (**C)** proportion of selected substitutions that were beneficial (*α*) in simulated non-coding regions. Note *α* was not estimated for DFE-alpha. All inferences were performed with 100k selected sites. In all panels, the error bars denote the standard deviation estimated from 10 independent replicates.

**Supplementary Figure 6:** Inference of the DFE and *α* under variation in ancestral population size. DFE inference was performed on an ingroup population (*N_cur_* = 1,000,000), which split from an outgroup population (*N_out_* = 1,500,000) 25*N_cur_* generations ago. The ancestral population size (*N_anc_*) was varied from 500,000 to 2,000,000. The DFE is shown in terms of the proportion of mutations in effectively neutral (*f*_0_), weakly deleterious (*f*_1_), moderately deleterious (*f*_2_) and strongly deleterious (*f*_3_) classes of mutations. For *α*, three simulated values are shown: the branch-specific proportion of adaptive substitutions in the ingroup (*α*_mel_), the branch-specific value in the outgroup (*α*_sim_), and the pooled value among substitutions accumulated on both branches (*α*_pairwise_). Error bars for DFE-alpha and GRAPES estimates denote the standard deviation across 10 simulation replicates.

**Supplementary Figure 7:** Inference of the distribution of fitness effects using the folded SFS as input for coding, high-confidence regulatory regions supported by empirical evidence from the REDfly database, and conserved non-coding regions. Left panels represent the West population and right panels represent the East population. The DFE is shown in terms of the proportion of mutations in effectively neutral (*f*_0_), weakly deleterious (*f*_1_), moderately deleterious (*f*_2_) and strongly deleterious (*f*_3_) classes of mutations, along with the proportion of substitutions that are beneficial (*α*). fastDFE *α* predictions were excluded due to their inaccuracy in the benchmarking simulations.

**Supplementary Figure 8:** Country-specific inferred distributions of fitness effects for high-confidence non-coding regulatory regions. Results are shown separately for Cameroon, Ghana, Gabon, and Rwanda using folded site-frequency spectra. The DFE is shown in terms of the proportion of mutations in effectively neutral (*f*_0_), weakly deleterious (*f*_1_), moderately deleterious (*f*_2_) and strongly deleterious (*f*_3_) classes of mutations.

**Supplementary Figure 9:** Inference of the distribution of fitness effects using the folded SFS as input for coding, low-confidence regulatory regions supported by computational evidence from FlyBase and OregAnno. Low-confidence regions overlapping phastCons elements were not included in this analysis. The DFE is shown in terms of the proportion of mutations in effectively neutral (*f*_0_), weakly deleterious (*f*_1_), moderately deleterious (*f*_2_) and strongly deleterious (*f*_3_) classes of mutations, along with the proportion of substitutions that are beneficial (*α*). fastDFE *α* predictions were excluded due to their inaccuracy in the benchmarking simulations.

**Supplementary Figure 10:** Inference of the distribution of fitness effects with and without masking sites within repeats for (**A**) transcription factor binding sites, (**B**) promoters, and (**C**) UTRs. The DFE is shown in terms of the proportion of mutations in effectively neutral (*f*_0_), weakly deleterious (*f*_1_), moderately deleterious (*f*_2_) and strongly deleterious (*f*_3_) classes of mutations, along with the proportion of substitutions that are beneficial (*α*). fastDFE *α* predictions were excluded due to their inaccuracy in the benchmarking simulations.

**Supplementary Figure 11:** Correlation between *B* inferred by Bvalcalc and nucleotide diversity (*π*) in the South population. Pearson’s *R*² and Spearman’s *ρ* were calculated from non-overlapping 1 Mb windows. Left: *B* maps computed using DFEs inferred from coding sites only. Right: *B* maps computed using DFEs inferred from coding sites, UTRs, and conserved non-coding elements.

**Supplementary Figure 12:** Sensitivity of neutral-site properties and DFE inference to per-site phastCons filtering. **(A)** Distribution of per-site phastCons scores at 0-fold coding sites and non-coding sites. “Other non-coding” represents the combination of high- and low-confidence annotations (not UTRs); phastCons-flanking neutral sites are overlaid in green. **(B)** Distribution of divergence across non-overlapping 100-kb windows containing at least 500 retained phastCons-flanking neutral sites. Dashed vertical lines indicate median divergence with no upper phastCons cutoff or a cutoff of ≤0.05. **(C)** Fraction of candidate neutral sites retained, nucleotide diversity (*π*), and divergence in the South population across maximum per-site phastCons-score cutoffs. The vertical dotted line marks the chosen cutoff of 0.1, and horizontal dashed lines show synonymous-site *π* and divergence. **(D)** Discretized DFE inferred for conserved non-coding regions using neutral sites filtered at the chosen cutoff of phastCons ≤0.10 or a stricter cutoff of phastCons ≤0.05.

**Supplementary Table 1:** Pairwise patterns of population differentiation (*F_ST_*) in fourfold degenerate sites, using the Weir-Cockerheim method implemented in scikit-allel. The number of individuals used for each country is shown on the corresponding diagonal. Pairwise *F_ST_* estimates among populations, country-level subsets, and inversion genotype classes. *F_ST_* was calculated at fourfold degenerate sites using the Weir-Cockerham estimator. Standard errors, 95% confidence intervals, and one-sided *p*-values for positive differentiation were estimated using a block-jackknife approach with non-overlapping 1 Mb genomic blocks. The “West” used in these comparisions excludes Ghana.

**Supplementary Table 2:** The nonoverlapping set of genomic sites used for DFE inference. For each class and subclass of sites we report the number of sites used for DFE inference, alongside the corresponding number of sites in the simulated genome and the percent of the simulated genome represented. “Neutral” subclasses (coding 4-fold sites and non-coding neutral sites) were used as putatively neutral reference sites; in particular, the non-coding neutral subclasses were drawn from the filtered set of intergenic neutral sites located within 5 kb of their paired selected site class (e.g., UTR, high-confidence regulatory regions, PhastCons elements, TFBS, promoters, enhancers), ensuring local matching of neutral and selected sites.

**Supplementary Table 3:** Summary of annotation counts and cumulative lengths of genomic elements used to define low-confidence regions restricted to *D. melanogaster* contigs 2L, 2R, 3L, and 3R. Totals reflect the number of annotated intervals and the summed interval lengths (bp) within each element–database category. Note these summaries are not mutually exclusive; annotations may overlap one another and may also overlap other feature classes (e.g., coding regions and *phastCons* elements).

**Supplementary Table 4:** DFE estimates for all three populations in all annotation types.

**Supplementary Table 5:** DFE estimates for four transcription quantiles of UniBind transcription factor binding sites in the South population.

