## Supplemental File for "The distribution of fitness effects of new mutations in regulatory regions of the *D. melanogaster* genome"

### SUPPLEMENTARY FIGURES AND TABLES

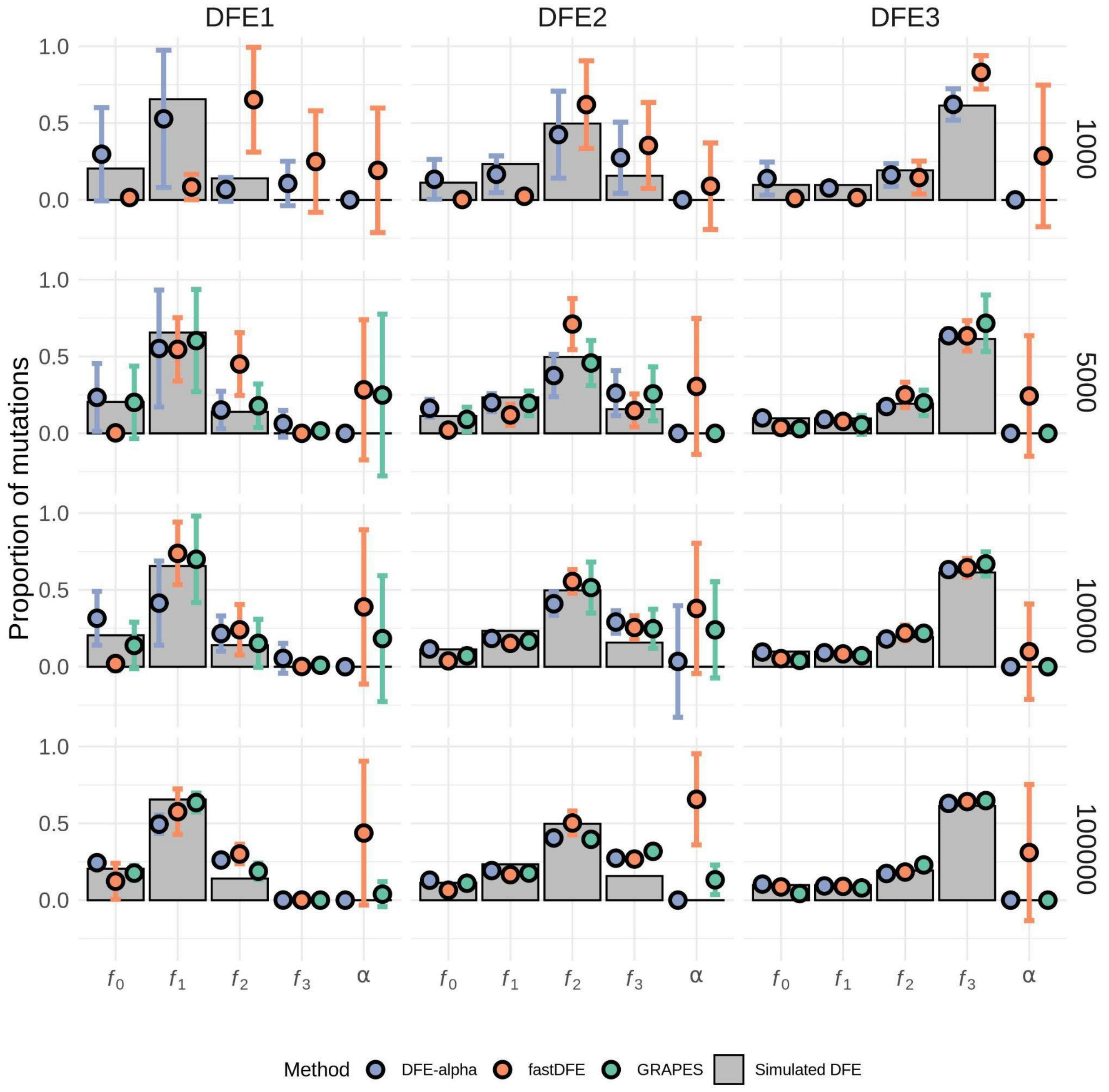


**Supplementary Figure 1:** The effects of the number of selected sites on the performance of *DFE-alpha*, *fastDFE*, and *GRAPES* in estimating the DFE of simulated high-confidence regulatory regions, when no beneficial mutations were simulated and the folded SFS was used as input. All inferences were performed with 200 individuals. Comparisons of the simulated and inferred DFE with a varying number of sites are shown in terms of the proportion of mutations in effectively neutral ($f_0$), weakly deleterious ($f_1$), moderately deleterious ($f_2$), and strongly deleterious ($f_3$) classes of mutations, along with the proportion of substitutions that are beneficial ($\alpha$). The simulated DFE is denoted with grey bars, and the inferred DFE is shown as circles. In all panels, the error bars denote the standard deviation of proportions estimated from 10 replicates where sites were drawn with replacement from the total number of simulated sites.

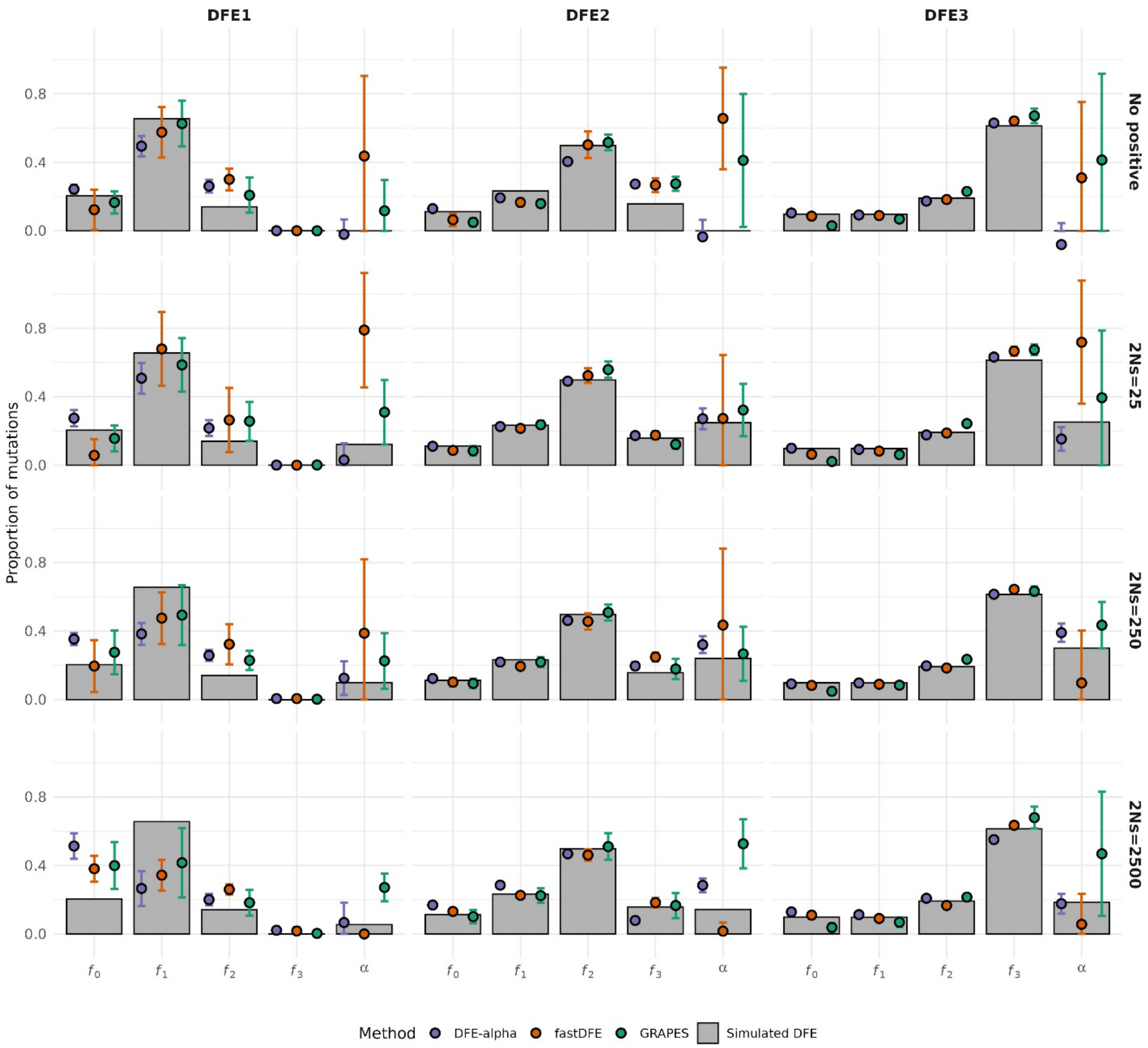


**Supplementary Figure 2:** The effects of strength and frequency of beneficial mutations on the performance of *DFE-alpha*, *fastDFE*, and *GRAPES* in estimating the DFE of simulated high-confidence regulatory regions, when no beneficial mutations were simulated and the folded SFS was used as input. All inferences were performed with 200 individuals. Comparisons of the simulated and inferred DFE with a varying number of sites are shown in terms of the proportion of mutations in effectively neutral ($f_0$), weakly deleterious ($f_1$), moderately deleterious ($f_2$), and strongly deleterious ($f_3$) classes of mutations, along with the proportion of substitutions that are beneficial ($\alpha$). The simulated DFE is denoted with grey bars, and the inferred DFE is shown as circles. In all panels, the error bars denote the standard deviation of proportions estimated from 10 replicates where sites were drawn with replacement from the total number of simulated sites.

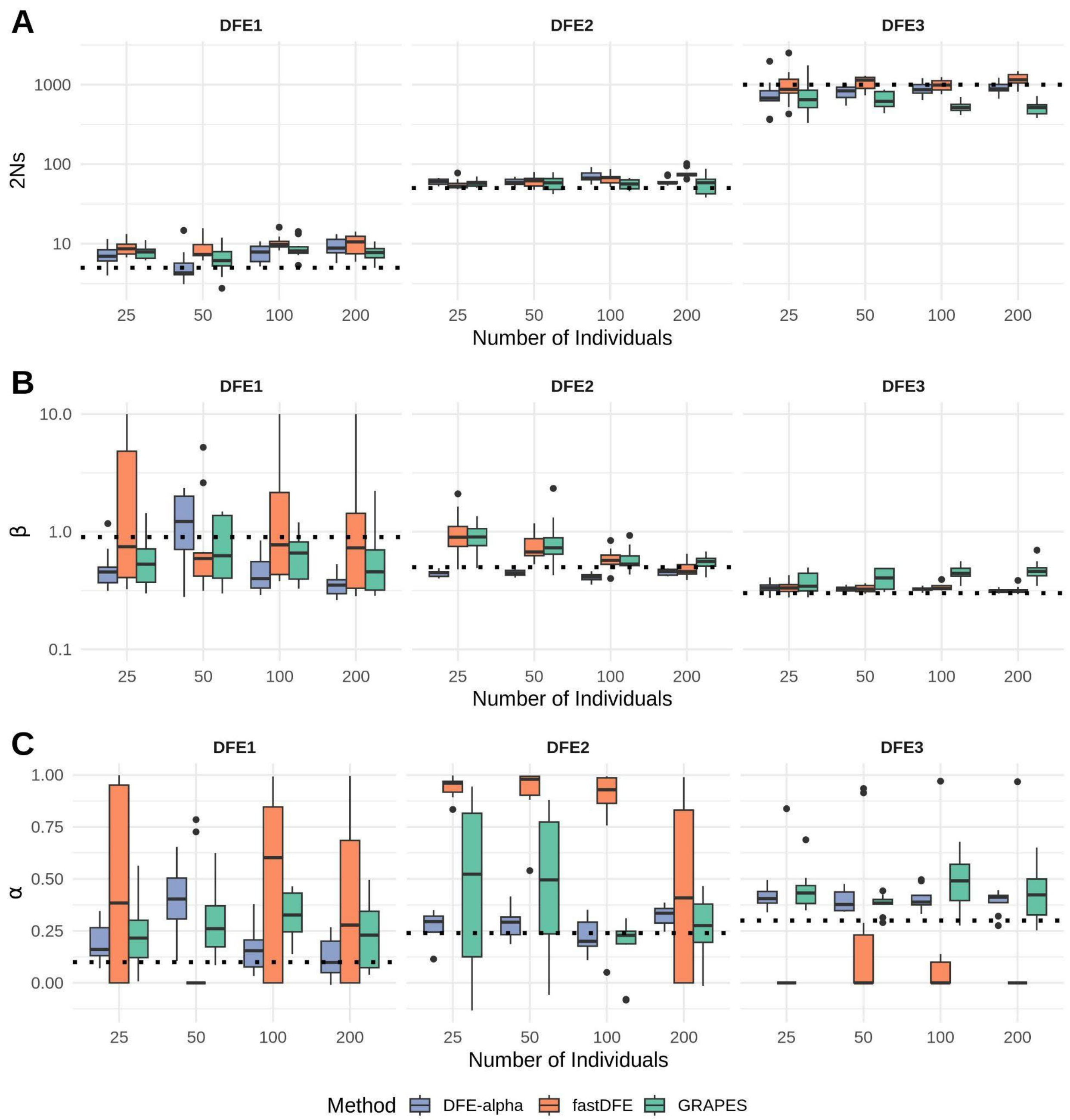


**Supplementary Figure 3:** Effects of sample size on the performance of *DFE-alpha*, *fastDFE*, and *GRAPES* on estimating the (**A**) mean ($\gamma$) and (**B**) shape ($\beta$) of the $\gamma$-distributed deleterious DFE, along with the (**C**) proportion of selected substitutions that were beneficial ($\alpha$) in simulated non-coding regions, using the folded SFS as input. Dotted lines represent the simulated parameters for $\gamma$ and $\beta$, and the true value of $\alpha$ or the full set of simulated high-confidence regions.

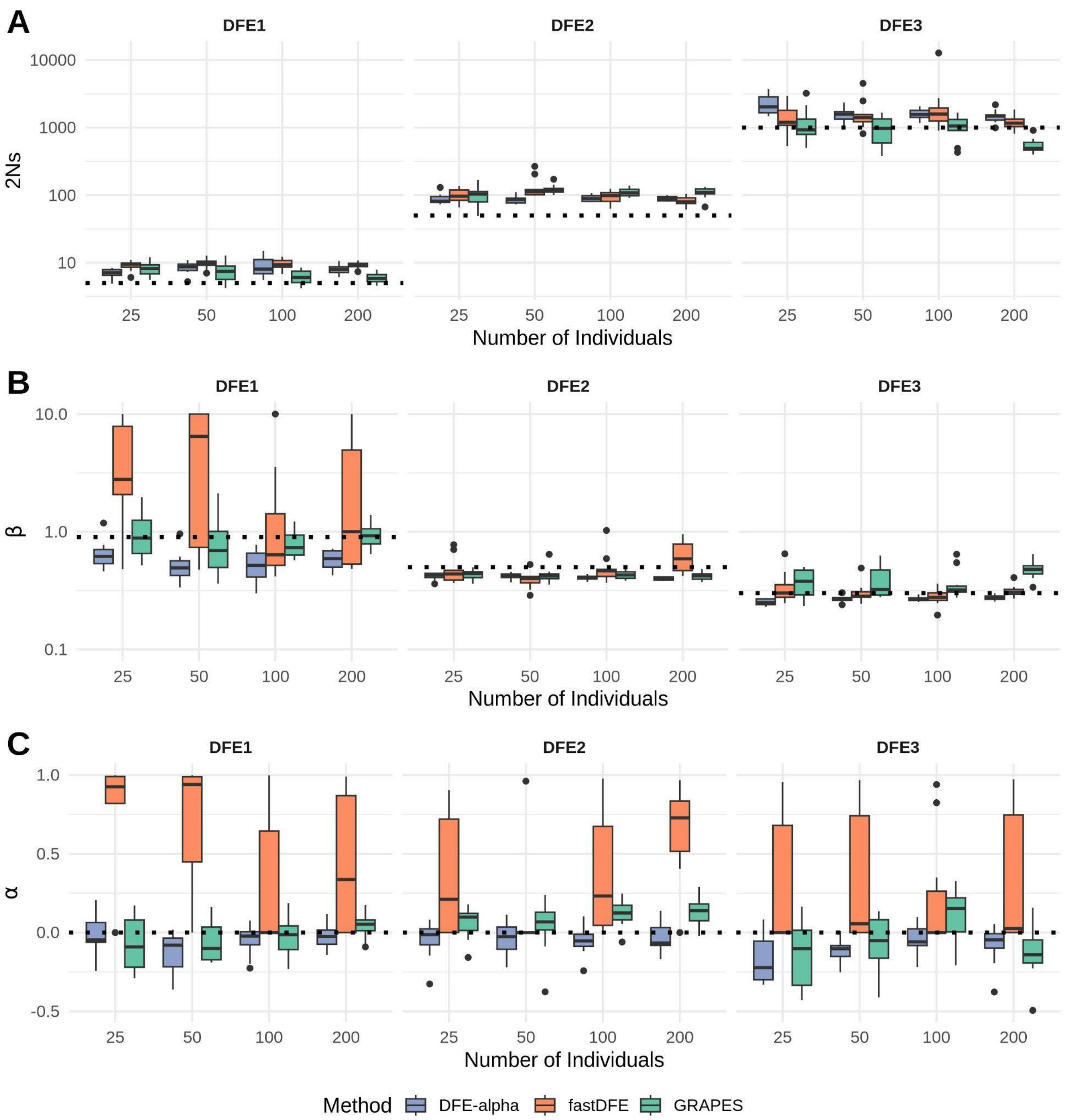


**Supplementary Figure 4:** Effects of varying the number of individuals on the performance of *DFE-alpha*, *fastDFE*, and *GRAPES* when no beneficial mutations were simulated and the folded SFS was used as input. Shown here are estimates of the (**A**) mean ($\gamma$) and (**B**) shape ($\beta$) of the $\gamma$-distributed deleterious DFE, along with the **(C)** proportion of selected substitutions that were beneficial ($\alpha$) in simulated non-coding regions. In all panels, the error bars denote the standard deviation estimated from 10 independent replicates.

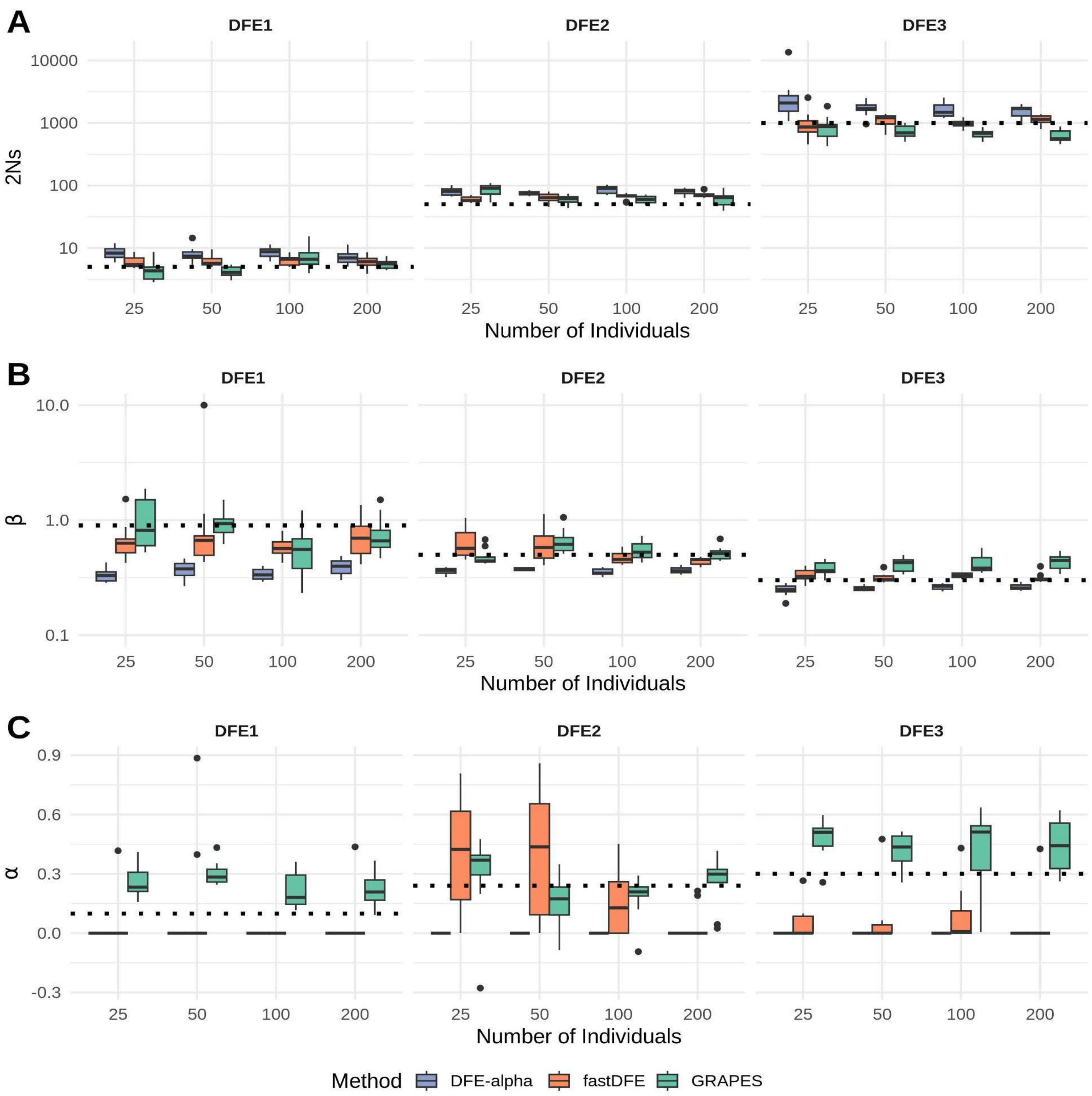


**Supplementary Figure 5:** Using the unfolded SFS, effects of varying the number of individuals on the performance of *DFE-alpha*, *fastDFE*, and *GRAPES* on estimating the (**A**) mean ($\gamma$) and (**B**) shape ($\beta$) of the $\gamma$-distributed deleterious DFE, along with the **(C)** proportion of selected substitutions that were beneficial ($\alpha$) in simulated non-coding regions. Note $\alpha$ was not estimated for DFE-alpha. All inferences were performed with 100k selected sites. In all panels, the error bars denote the standard deviation estimated from 10 independent replicates.

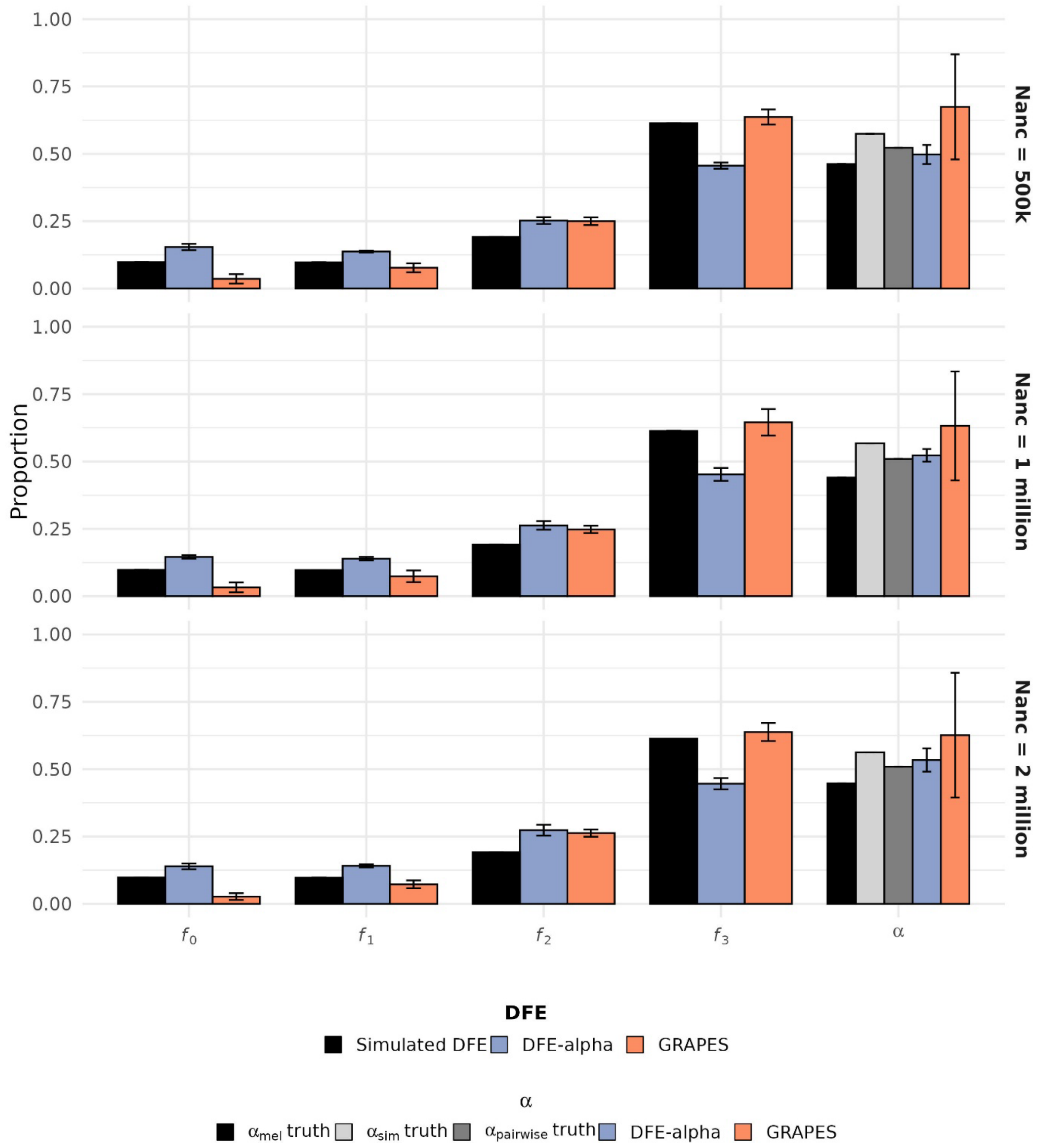


**Supplementary Figure 6:** Inference of the DFE and $\alpha$ under variation in ancestral population size. DFE inference was performed on an ingroup population ($N_{cur} = 1{,}000{,}000$), which split from an outgroup population ($N_{out} = 1{,}500{,}000$) $25N_{cur}$ generations ago. The ancestral population size ($N_{anc}$) was varied from 500,000 to 2,000,000. The DFE is shown in terms of the proportion of mutations in effectively neutral ($f_0$), weakly deleterious ($f_1$), moderately deleterious ($f_2$) and strongly deleterious ($f_3$) classes of mutations. For $\alpha$, three simulated values are shown: the branch-specific proportion of adaptive substitutions in the ingroup ($\alpha_{\text{mel}}$), the branch-specific value in the outgroup ($\alpha_{\text{sim}}$), and the pooled value among substitutions accumulated on both branches ($\alpha_{\text{pairwise}}$). Error bars for DFE-alpha and GRAPES estimates denote the standard deviation across 10 simulation replicates.

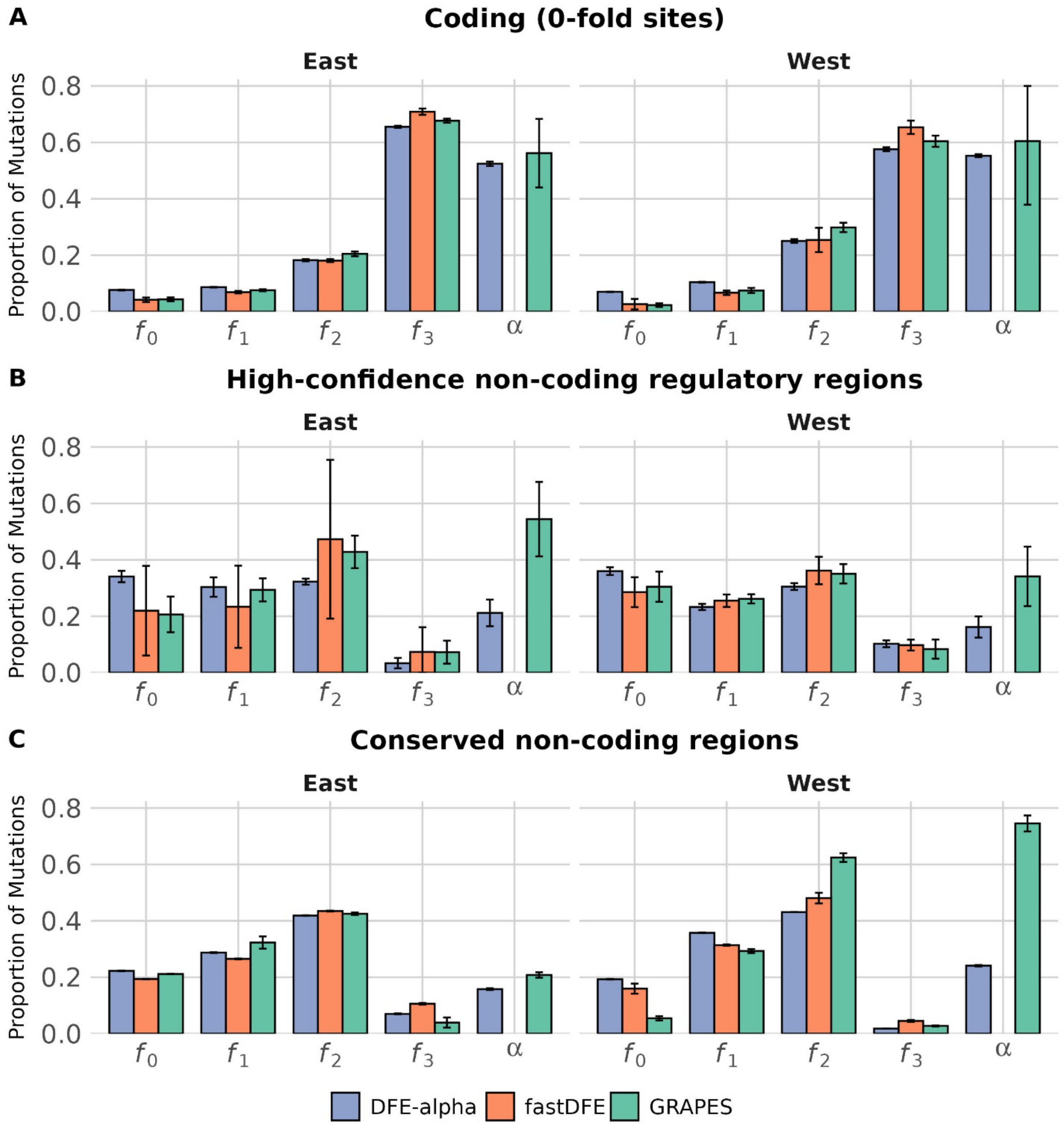


**Supplementary Figure 7:** Inference of the distribution of fitness effects using the folded SFS as input for coding, high-confidence regulatory regions supported by empirical evidence from the REDfly database, and conserved non-coding regions. Left panels represent the West population and right panels represent the East population. The DFE is shown in terms of the proportion of mutations in effectively neutral ($f_0$), weakly deleterious ($f_1$), moderately deleterious ($f_2$) and strongly deleterious ($f_3$) classes of mutations, along with the proportion of substitutions that are beneficial ($\alpha$). fastDFE $\alpha$ predictions were excluded due to their inaccuracy in the benchmarking simulations.

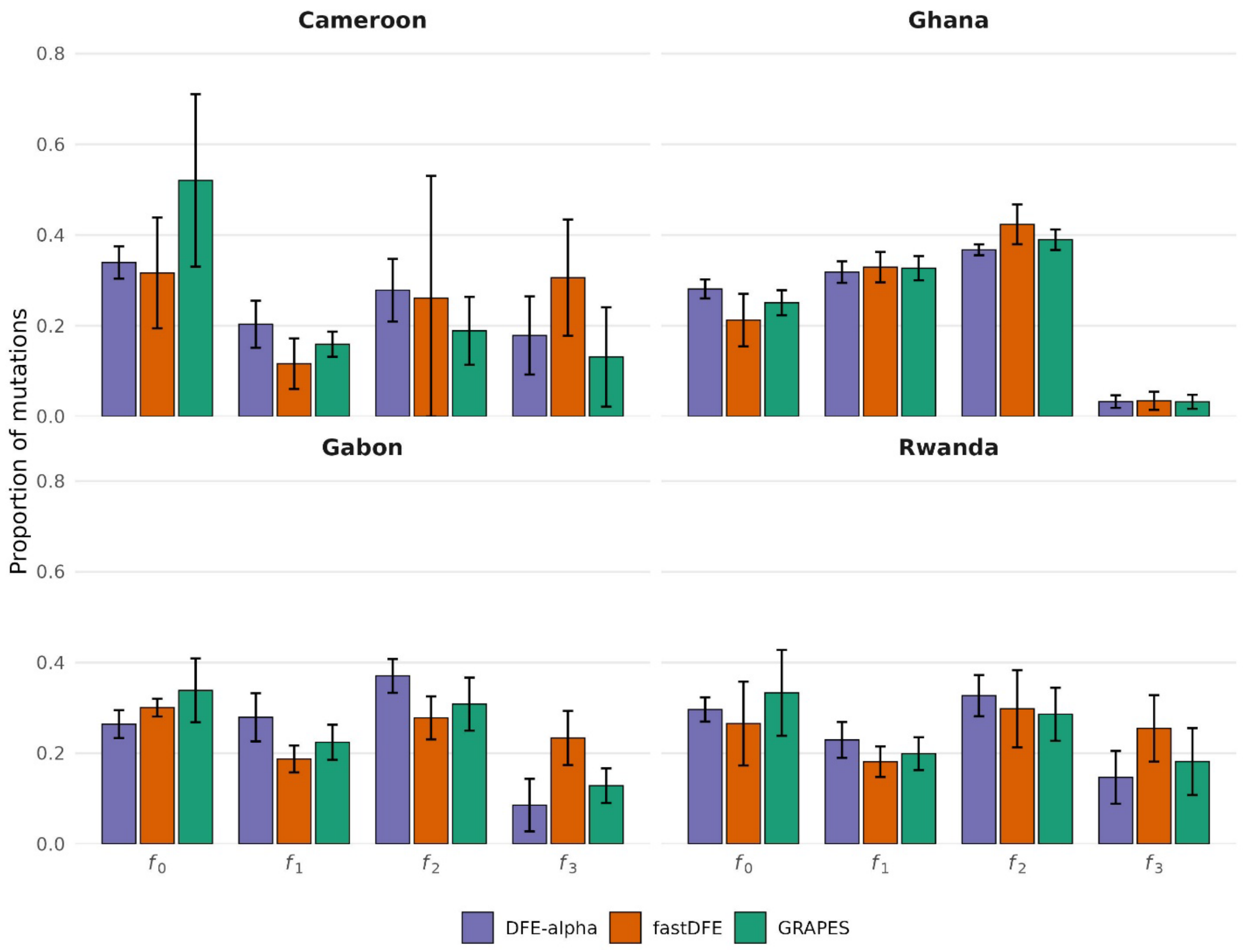


**Supplementary Figure 8:** Country-specific inferred distributions of fitness effects for high-confidence non-coding regulatory regions. Results are shown separately for Cameroon, Ghana, Gabon, and Rwanda using folded site-frequency spectra. The DFE is shown in terms of the proportion of mutations in effectively neutral ($f_0$), weakly deleterious ($f_1$), moderately deleterious ($f_2$) and strongly deleterious ($f_3$) classes of mutations.

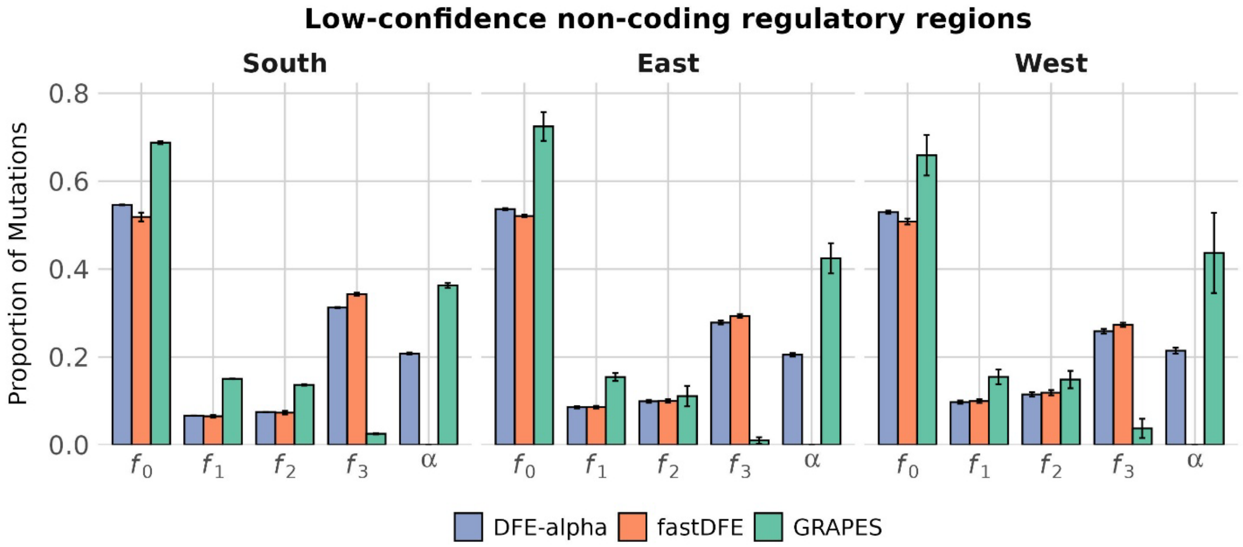


**Supplementary Figure 9:** Inference of the distribution of fitness effects using the folded SFS as input for coding, low-confidence regulatory regions supported by computational evidence from FlyBase and OregAnno. Low-confidence regions overlapping phastCons elements were not included in this analysis. The DFE is shown in terms of the proportion of mutations in effectively neutral ($f_0$), weakly deleterious ($f_1$), moderately deleterious ($f_2$) and strongly deleterious ($f_3$) classes of mutations, along with the proportion of substitutions that are beneficial ($\alpha$). fastDFE $\alpha$ predictions were excluded due to their inaccuracy in the benchmarking simulations.

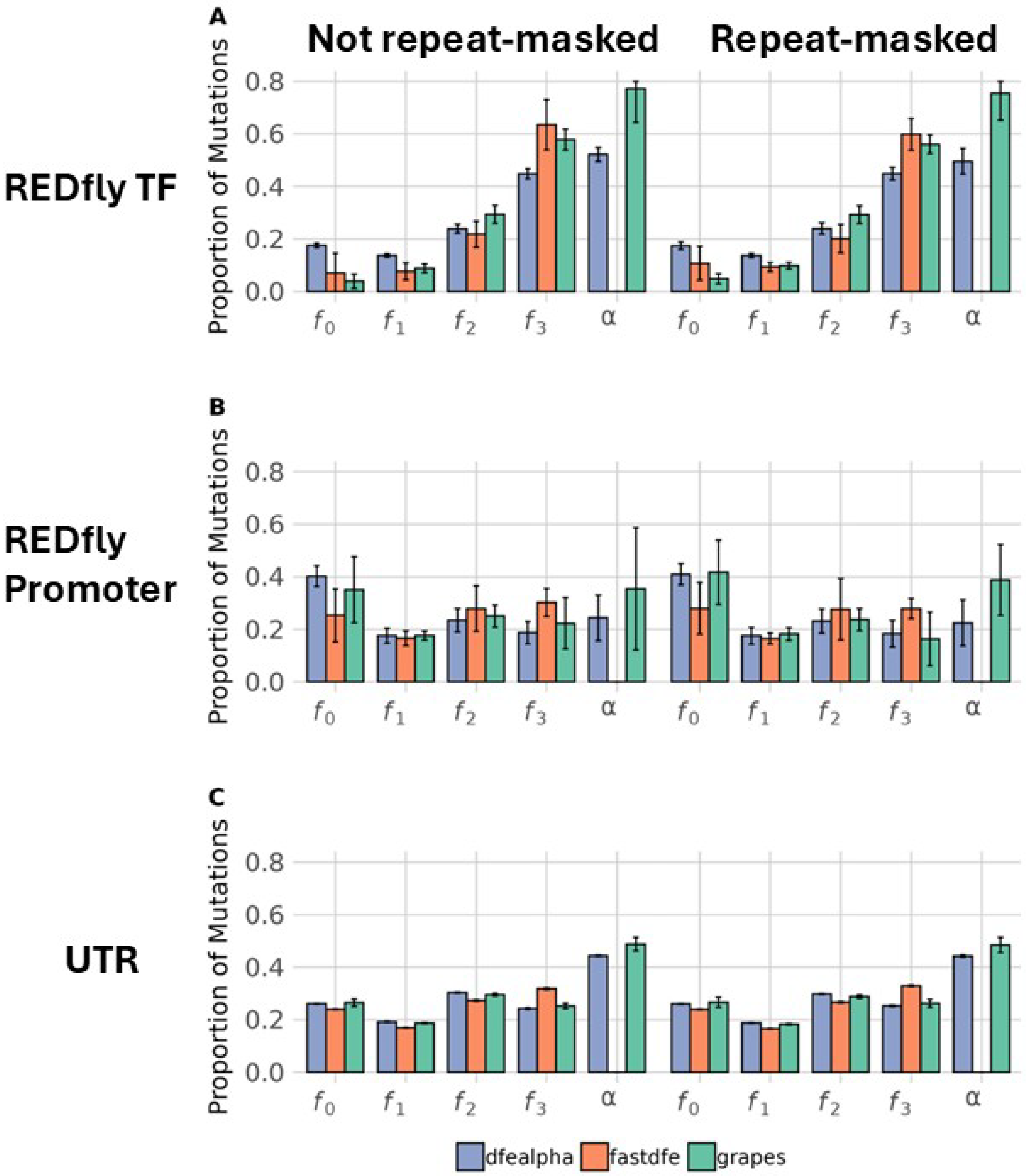


**Supplementary Figure 10:** Inference of the distribution of fitness effects with and without masking sites within repeats for (**A**) transcription factor binding sites, (**B**) promoters, and (**C**) UTRs. The DFE is shown in terms of the proportion of mutations in effectively neutral ($f_0$), weakly deleterious ($f_1$), moderately deleterious ($f_2$) and strongly deleterious ($f_3$) classes of mutations, along with the proportion of substitutions that are beneficial ($\alpha$). fastDFE $\alpha$ predictions were excluded due to their inaccuracy in the benchmarking simulations.

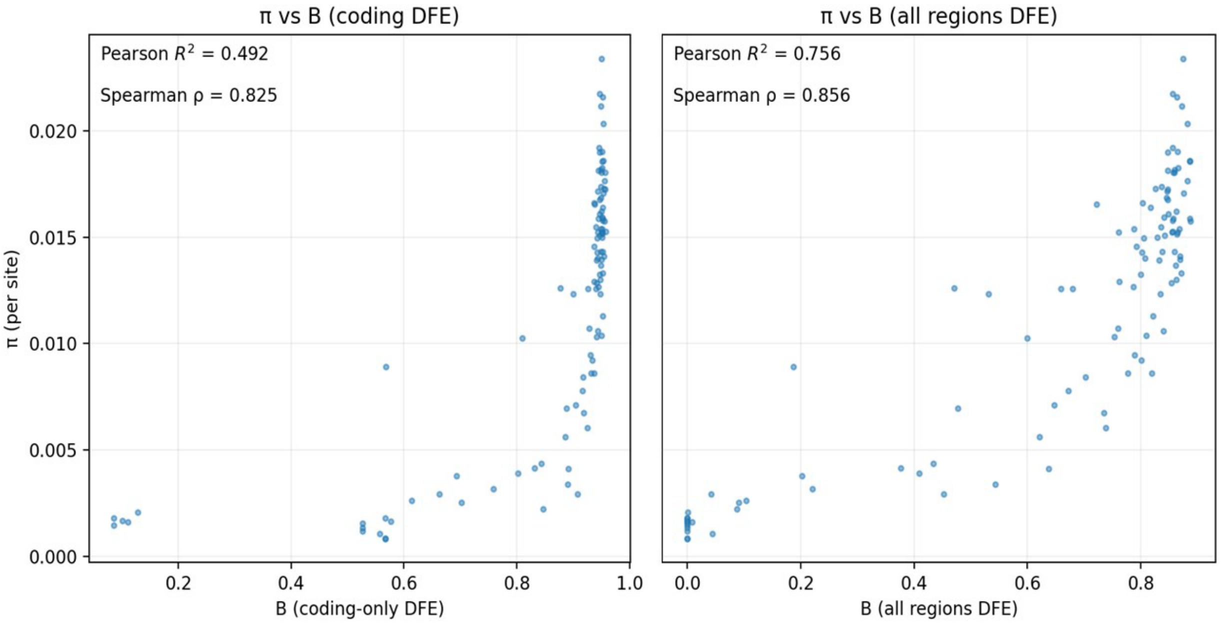


**Supplementary Figure 11:** Correlation between $B$ inferred by Bvalcalc and nucleotide diversity ($\pi$) in the South population. Pearson's $R^2$ and Spearman's $\rho$ were calculated from non-overlapping 1 Mb windows. Left: $B$ maps computed using DFEs inferred from coding sites only. Right: $B$ maps computed using DFEs inferred from coding sites, UTRs, and conserved non-coding elements.

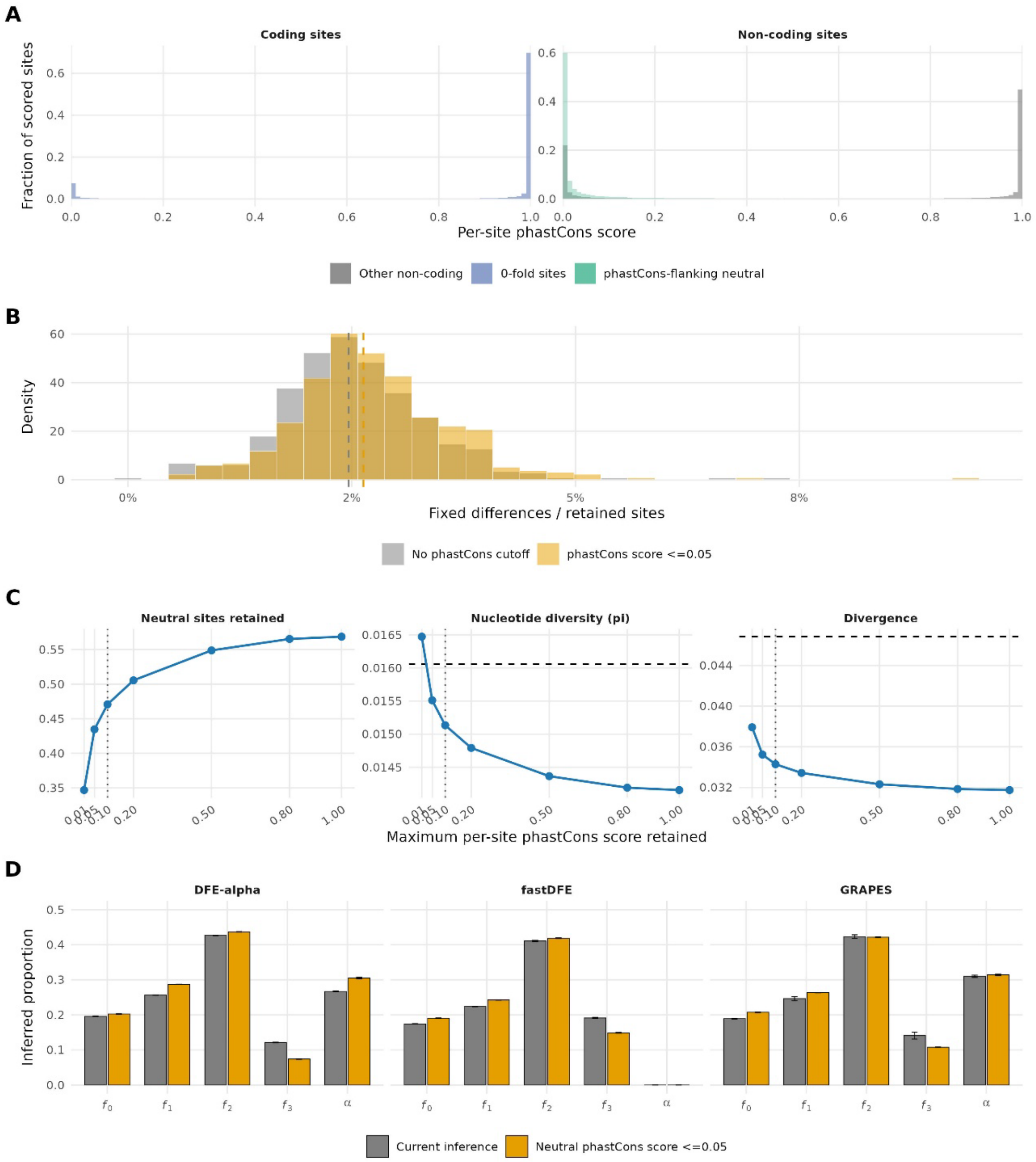


**Supplementary Figure 12:** Sensitivity of neutral-site properties and DFE inference to per-site phastCons filtering. **(A)** Distribution of per-site phastCons scores at 0-fold coding sites and non-coding sites. "Other non-coding" represents the combination of high- and low-confidence annotations (not UTRs); phastCons-flanking neutral sites are overlaid in green. **(B)** Distribution of divergence across non-overlapping 100-kb windows containing at least 500 retained phastCons-flanking neutral sites. Dashed vertical lines indicate median divergence with no upper phastCons cutoff or a cutoff of ≤0.05. **(C)** Fraction of candidate neutral sites retained, nucleotide diversity ($\pi$), and divergence in the South population across maximum per-site phastCons-score cutoffs. The vertical dotted line marks the chosen cutoff of 0.1, and horizontal dashed lines show synonymous-site $\pi$ and divergence. **(D)** Discretized DFE inferred for conserved non-coding regions using neutral sites filtered at the chosen cutoff of phastCons ≤0.10 or a stricter cutoff of phastCons ≤0.05.

**Supplementary Table 1:** Pairwise patterns of population differentiation ($F_{ST}$) in fourfold degenerate sites, using the Weir-Cockerheim method implemented in scikit-allel. The number of individuals used for each country is shown on the corresponding diagonal. Pairwise $F_{ST}$ estimates among populations, country-level subsets, and inversion genotype classes. $F_{ST}$ was calculated at fourfold degenerate sites using the Weir-Cockerham estimator. Standard errors, 95% confidence intervals, and one-sided *p*-values for positive differentiation were estimated using a block-jackknife approach with non-overlapping 1 Mb genomic blocks. The "West" used in these comparisions excludes Ghana.

| Comparison | $F_{ST}$ mean | 95% CI | *P* value |
|---|---|---|---|
| South vs West | 0.0975 | 0.0833-0.1116 | 9.39e-42 |
| South vs East | 0.0781 | 0.0630-0.0931 | 1.36e-24 |
| East vs West | 0.0391 | 0.0327-0.0456 | 7.63e-33 |
| Uganda vs Rwanda | -0.0184 | -0.0240--0.0128 | 1.0 |
| Ghana vs Cameroon | 0.0335 | 0.0261-0.0409 | 4.36e-19 |
| Ghana vs Gabon | 0.0224 | 0.0167-0.0281 | 9.54e-15 |
| Ghana vs Guinea | 0.0285 | 0.0239-0.0332 | 1.49e-33 |
| Ghana vs Nigeria | 0.0223 | 0.0182-0.0264 | 1.08e-26 |
| East In(3R)P_e | -0.0188 | -0.0230--0.0145 | 1.0 |
| West In(2L)t | 0.0028 | -0.0052-0.0109 | 0.246 |
| West In(2R)NS | -0.0805 | -0.0887--0.0724 | 1.0 |
| West In(3L)P | -0.0236 | -0.0326--0.0147 | 1.0 |
| West In(3R)K | -0.0033 | -0.0067-0.0001 | 0.973 |
| West In(3R)P_e | -0.0146 | -0.0183-0.0109 | 1.0 |

**Supplementary Table 2:** The nonoverlapping set of genomic sites used for DFE inference. For each class and subclass of sites we report the number of sites used for DFE inference, alongside the corresponding number of sites in the simulated genome and the percent of the simulated genome represented. "Neutral" subclasses (coding 4-fold sites and non-coding neutral sites) were used as putatively neutral reference sites; in particular, the non-coding neutral subclasses were drawn from the filtered set of intergenic neutral sites located within 5 kb of their paired selected site class (e.g., UTR, high-confidence regulatory regions, PhastCons elements, TFBS, promoters, enhancers), ensuring local matching of neutral and selected sites.

| Site class | Site subclass | Number of sites used for inference | Simulated number of sites | % of simulated genome |
|---|---|---|---|---|
| Exons | | | 16,873,374 | 17.45 |
| | Selected coding (0-fold) | 3,515,727 | | |
| | Neutral coding (4-fold) | 832,622 | | |
| | Untranslated region (5' & 3') | 3,507,752 | 7,529,622 | 7.79 |
| High-confidence regulatory region | | 82,882 | 216,601 | 0.22 |
| | Transcription factor binding sites | 14,022 | | |
| | Promoters | 18,592 | | |
| | Enhancers | 50,261 | | |
| Low-confidence regulatory region | | | 60,672,393 | 62.74 |

| | | | | |
|---|---|---|---|---|
| | PhastCons elements | 16,642,807 | | |
| Non-coding neutral | | | 11407698 | 11.8 |
| | Neutral flanking, UTR | 840,841 | | |
| | Neutral flanking, high confidence | 64,627 | | |
| | Neutral flanking, phastCons elements | 1,364,872 | | |
| | Neutral flanking, TFBS | 25,111 | | |
| | Neutral flanking, promoter | 8,612 | | |
| | Neutral flanking, enhancer | 4,735 | | |

**Supplementary Table 3:** Summary of annotation counts and cumulative lengths of genomic elements used to define low-confidence regions restricted to *D. melanogaster* contigs 2L, 2R, 3L, and 3R. Totals reflect the number of annotated intervals and the summed interval lengths (bp) within each element–database category. Note these summaries are not mutually exclusive; annotations may overlap one another and may also overlap other feature classes (e.g., coding regions and *phastCons* elements).

| element | Database | Count | Total Length |
|---|---|---:|---:|
| Transcription Factor Binding Site | FlyBase | 194019 | 145399432 |
| Regulatory Region | FlyBase | 17004 | 8955659 |
| deletion | FlyBase | 4957 | 6877154 |
| ncRNA | FlyBase | 2580 | 4925839 |
| RNAi_reagent | FlyBase | 5881 | 1836864 |
| delins | FlyBase | 1754 | 1312544 |
| Regulatory Region | ORegAnno | 560 | 960657 |
| TSS | FlyBase | 6995 | 272855 |
| insulator | FlyBase | 6199 | 61990 |
| sgRNA | FlyBase | 1874 | 35608 |
| Transcription Factor Binding Site | ORegAnno | 1653 | 32537 |
| snoRNA | FlyBase | 238 | 27986 |
| tRNA | FlyBase | 266 | 19884 |
| mature_protein_region | FlyBase | 7 | 18688 |
| pre_miRNA | FlyBase | 200 | 18019 |
| insertion | FlyBase | 221 | 15306 |
| rRNA | FlyBase | 96 | 12862 |
| miRNA | FlyBase | 368 | 7834 |
| miRNA_primary_transcript | miRBase | 71 | 6404 |
| snRNA | FlyBase | 30 | 5069 |

| element | Database | Count | Total Length |
|---|---|---|---|
| miRNA | miRBase | 115 | 2429 |
| insertion_site | FlyBase | 10 | 1948 |
| miRNA Binding Site | ORegAnno | 64 | 1466 |
| protein_binding_site | FlyBase | 27 | 742 |

**Supplementary Table 4:** DFE estimates for all three populations in all annotation types.

| Method | Pop. | γ mean | γ SD | β mean | β SD | α mean | α SD | Annotation |
|---|---|---|---|---|---|---|---|---|
| DFE-alpha | South | 811.4 | 26.991 | 0.347 | 0.005 | 0.536 | 0.008 | Coding (0-fold sites) |
| fastDFE | South | -1,983.7 | 93.357 | 0.312 | 0.004 | 0.000 | 0.000 | Coding (0-fold sites) |
| GRAPES | South | 993.5 | 85.159 | 0.353 | 0.018 | 0.557 | 0.030 | Coding (0-fold sites) |
| DFE-alpha | East | 2,293.5 | 198.393 | 0.323 | 0.021 | 0.509 | 0.024 | Coding (0-fold sites) |
| fastDFE | East | -2,023.7 | 246.231 | 0.304 | 0.008 | 0.000 | 0.000 | Coding (0-fold sites) |
| GRAPES | East | 1,316.7 | 143.755 | 0.307 | 0.008 | 0.495 | 0.012 | Coding (0-fold sites) |
| DFE-alpha | West | 777.8 | 89.243 | 0.332 | 0.009 | 0.470 | 0.012 | Coding (0-fold sites) |
| fastDFE | West | -817.0 | 86.834 | 0.355 | 0.011 | 0.000 | 0.000 | Coding (0-fold sites) |
| GRAPES | West | 555.3 | 59.007 | 0.360 | 0.011 | 0.508 | 0.011 | Coding (0-fold sites) |
| DFE-alpha | South | 47.2 | 6.861 | 0.210 | 0.019 | 0.189 | 0.040 | High-confidence non-coding regulatory regions |
| fastDFE | South | -72.0 | 17.994 | 0.251 | 0.048 | 0.696 | 0.384 | High-confidence non-coding regulatory regions |
| GRAPES | South | -79.5 | 24.523 | 0.309 | 0.063 | 0.520 | 0.093 | High-confidence non-coding regulatory regions |
| DFE-alpha | East | -70.7 | 14.997 | 0.295 | 0.045 | 0.34.2 | 0.041 | High-confidence non-coding regulatory regions |

| Method | Pop. | γ mean | γ SD | β mean | β SD | α mean | α SD | Annotation |
|---|---|---|---|---|---|---|---|---|
| fastDFE | East | -71.7 | 22.838 | 0.246 | 0.047 | 0.089 | 0.281 | High-confidence non-coding regulatory regions |
| GRAPES | East | -41.8 | 10.130 | 0.227 | 0.031 | 0.297 | 0.036 | High-confidence non-coding regulatory regions |
| DFE-alpha | West | -99.1 | 59.678 | 0.234 | 0.054 | 0.257 | 0.060 | High-confidence non-coding regulatory regions |
| fastDFE | West | -95.4 | 64.028 | 0.315 | 0.279 | 0.308 | 0.402 | High-confidence non-coding regulatory regions |
| GRAPES | West | -54.6 | 14.834 | 0.236 | 0.093 | 0.371 | 0.158 | High-confidence non-coding regulatory regions |
| DFE-alpha | South | -37.9 | 0.316 | 0.371 | 0.003 | 0.269 | 0.003 | phastCons |
| fastDFE | South | -59.8 | 0.632 | 0.367 | 0.005 | 0.000 | 0.000 | phastCons |
| GRAPES | South | -45.3 | 2.584 | 0.371 | 0.007 | 0.311 | 0.003 | phastCons |
| DFE-alpha | East | -82.4 | 0.516 | 0.420 | 0.000 | 0.280 | 0.000 | phastCons |
| fastDFE | East | -49.7 | 0.483 | 0.351 | 0.003 | 0.000 | 0.000 | phastCons |
| GRAPES | East | -28.8 | 7.177 | 0.391 | 0.031 | 0.232 | 0.013 | phastCons |
| DFE-alpha | West | -22.7 | 0.483 | 0.442 | 0.006 | 0.262 | 0.004 | phastCons |
| fastDFE | West | -30.4 | 0.516 | 0.446 | 0.005 | 0.001 | 0.003 | phastCons |
| GRAPES | West | -21.4 | 4.427 | 0.478 | 0.043 | 0.281 | 0.014 | phastCons |
| DFE-alpha | South | -104.4 | 83.520 | 0.161 | 0.034 | 0.245 | 0.085 | REDfly promoter |
| fastDFE | South | -210.6 | 153.739 | 0.245 | 0.099 | 0.770 | 0.410 | REDfly promoter |
| GRAPES | South | -105.5 | 60.748 | 0.193 | 0.051 | 0.354 | 0.233 | REDfly promoter |
| DFE-alpha | East | -42.6 | 36.540 | 0.328 | 0.202 | 0.401 | 0.086 | REDfly promoter |
| fastDFE | East | -68.6 | 94.129 | 0.490 | 0.554 | 0.220 | 0.377 | REDfly promoter |

| Method | Pop. | γ mean | γ SD | β mean | β SD | α mean | α SD | Annotation |
|---|---|---|---|---|---|---|---|---|
| GRAPES | East | -37.4 | 40.247 | 0.306 | 0.144 | 0.484 | 0.150 | REDfly promoter |
| DFE-alpha | West | -218.4 | 191.28 | 0.188 | 0.092 | 0.331 | 0.116 | REDfly promoter |
| fastDFE | West | -20,042 | 42,141 | 6.001 | 4.610 | 0.896 | 0.315 | REDfly promoter |
| GRAPES | West | -113.6 | 90.818 | 0.285 | 0.127 | 0.566 | 0.182 | REDfly promoter |
| DFE-alpha | South | -38.9 | 14.896 | 0.162 | 0.030 | -0.070 | 0.073 | REDfly enhancer |
| fastDFE | South | -74.6 | 36.809 | 0.179 | 0.053 | 0.069 | 0.151 | REDfly enhancer |
| GRAPES | South | -43.4 | 22.476 | 0.225 | 0.033 | 0.259 | 0.159 | REDfly enhancer |
| DFE-alpha | East | -10,662 | 19,009 | 0.217 | 0.137 | 0.145 | 0.195 | REDfly enhancer |
| fastDFE | East | -20,023 | 42,152 | 9.125 | 1.872 | 1.000 | 0.000 | REDfly enhancer |
| GRAPES | East | -109.4 | 115.94 | 0.221 | 0.070 | 0.239 | 0.296 | REDfly enhancer |
| DFE-alpha | West | -48.5 | 19.010 | 0.414 | 0.172 | 0.260 | 0.122 | REDfly enhancer |
| fastDFE | West | -14.8 | 5.116 | 7.317 | 4.335 | 0.971 | 0.066 | REDfly enhancer |
| GRAPES | West | -19.2 | 11.811 | 0.548 | 0.251 | 0.512 | 0.235 | REDfly enhancer |
| DFE-alpha | South | -93.3 | 2.263 | 0.240 | 0.000 | 0.443 | 0.005 | UTR |
| fastDFE | South | -161.4 | 4.993 | 0.231 | 0.003 | 0.000 | 0.000 | UTR |
| GRAPES | South | -101.6 | 5.358 | 0.236 | 0.010 | 0.489 | 0.025 | UTR |
| DFE-alpha | East | -160.2 | 5.514 | 0.280 | 0.008 | 0.459 | 0.007 | UTR |
| fastDFE | East | -125.4 | 8.527 | 0.233 | 0.007 | 0.000 | 0.000 | UTR |
| GRAPES | East | -71.4 | 11.098 | 0.221 | 0.009 | 0.418 | 0.006 | UTR |
| DFE-alpha | West | -47.9 | 2.558 | 0.282 | 0.006 | 0.436 | 0.007 | UTR |
| fastDFE | West | -54.8 | 3.225 | 0.321 | 0.018 | 0.409 | 0.285 | UTR |
| GRAPES | West | -43.6 | 2.221 | 0.293 | 0.014 | 0.459 | 0.026 | UTR |
| DFE-alpha | South | -387.4 | 86.982 | 0.251 | 0.014 | 0.523 | 0.025 | REDfly TFBS |
| fastDFE | South | -1,165.3 | 2,034.613 | 0.506 | 0.244 | 0.714 | 0.381 | REDfly TFBS |

| Method | Pop. | γ mean | γ SD | β mean | β SD | α mean | α SD | Annotation |
|---|---|---|---|---|---|---|---|---|
| GRAPES | South | -308.2 | 94.788 | 0.565 | 0.111 | 0.772 | 0.128 | REDfly TFBS |
| DFE-alpha | East | -110.5 | 19.196 | 0.584 | 0.076 | 0.695 | 0.046 | REDfly TFBS |
| fastDFE | East | -52.5 | 17.367 | 1.416 | 2.685 | 0.205 | 0.342 | REDfly TFBS |
| GRAPES | East | -41.6 | 14.385 | 0.659 | 0.265 | 0.700 | 0.103 | REDfly TFBS |
| DFE-alpha | West | -233.7 | 51.130 | 0.457 | 0.063 | 0.662 | 0.037 | REDfly TFBS |
| fastDFE | West | -63.6 | 31.210 | 2.057 | 2.844 | 0.987 | 0.028 | REDfly TFBS |
| GRAPES | West | -81.8 | 39.988 | 10.672 | 31.389 | 0.718 | 0.189 | REDfly TFBS |
| DFE-alpha | South | -39.1 | 1.595 | 0.322 | 0.009 | 0.368 | 0.016 | UniBind TFBS |
| fastDFE | South | -60.6 | 3.169 | 0.319 | 0.012 | 0.000 | 0.000 | UniBind TFBS |
| GRAPES | South | -45.0 | 3.559 | 0.321 | 0.014 | 0.412 | 0.018 | UniBind TFBS |
| DFE-alpha | East | -65.7 | 3.561 | 0.415 | 0.020 | 0.418 | 0.015 | UniBind TFBS |
| fastDFE | East | -39.2 | 3.490 | 0.380 | 0.041 | 0.443 | 0.269 | UniBind TFBS |
| GRAPES | East | -33.0 | 2.309 | 0.335 | 0.018 | 0.361 | 0.017 | UniBind TFBS |
| DFE-alpha | West | -19.5 | 2.506 | 0.440 | 0.037 | 0.410 | 0.029 | UniBind TFBS |
| fastDFE | West | -21.6 | 2.066 | 1.168 | 0.305 | 0.954 | 0.026 | UniBind TFBS |
| GRAPES | West | -24.4 | 4.169 | 0.657 | 0.102 | 0.623 | 0.165 | UniBind TFBS |

**Supplementary Table 5:** DFE estimates for four transcription quantiles of UniBind transcription factor binding sites in the South population.

| Caller | γ mean | γ SD | β mean | β SD | α mean | α SD | Quantile |
|---|---|---|---|---|---|---|---|
| DFE-alpha | -30.5 | 2.718 | 0.357 | 0.021 | 0.364 | 0.037 | Q1 |
| GRAPES | -24.9 | 5.301 | 0.397 | 0.038 | 0.405 | 0.030 | |
| fastDFE | -47.0 | 6.799 | 0.344 | 0.020 | 0.000 | 0.000 | |
| DFE-alpha | -30.0 | 3.590 | 0.339 | 0.023 | 0.360 | 0.040 | Q2 |
| GRAPES | -47.6 | 13.418 | 0.437 | 0.087 | 0.464 | 0.218 | |
| fastDFE | -50.3 | 13.581 | 0.429 | 0.097 | 0.803 | 0.273 | |
| DFE-alpha | -24.6 | 2.503 | 0.369 | 0.034 | 0.325 | 0.040 | Q3 |
| GRAPES | -29.2 | 7.913 | 0.360 | 0.047 | 0.323 | 0.048 | |
| fastDFE | -44.6 | 7.633 | 0.331 | 0.034 | 0.000 | 0.000 | |
| DFE-alpha | -57.1 | 7.445 | 0.260 | 0.014 | 0.429 | 0.027 | Q4 |
| GRAPES | -68.3 | 12.988 | 0.356 | 0.041 | 0.396 | 0.099 | |
| fastDFE | -98.6 | 14.230 | 0.345 | 0.160 | 0.537 | 0.425 | |
